# Off-manifold coding in visual cortex revealed by sleep

**DOI:** 10.1101/2022.06.10.495710

**Authors:** Eliezyer Fermino de Oliveira, Soyoun Kim, Tian Season Qiu, Adrien Peyrache, Renata Batista-Brito, Lucas Sjulson

## Abstract

Low-dimensional neural manifolds are controversial in part because it is unclear how to reconcile them with high-dimensional representations observed in areas such as primary visual cortex (V1). We addressed this by recording neuronal activity in V1 during slow-wave sleep, enabling us to identify internally-generated low-dimensional manifold structure and evaluate its role during visual processing. We found that movements and visual stimuli were both encoded in the “on-manifold” subspace preserved during sleep. However, only stimuli were encoded in the “off-manifold” subspace, which contains activity patterns that are less likely than chance to occur spontaneously during sleep. This off-manifold activity comprises sparse firing in neurons with the strongest low-dimensional modulation by movement, which paradoxically prevents movement-evoked activity from interfering with stimulus representations. These results reveal an unexpected link between low-dimensional dynamics and sparse coding, which together create a protected off-manifold coding space keeping high-dimensional representations separable from movement-evoked activity.

## Introduction

Recent studies using large-scale neuronal recordings have reported that neural activity contains brain-wide low-dimensional representations of movements and internal states (1–4), even in sensory areas such as primary visual cortex (V1). Many studies have also suggested that the brain operates in a low-dimensional dynamical regime, with activity constrained to a neural “manifold,” or low-dimensional subspace (5–10). However, high-dimensional representations of sensory stimuli and behavioral variables have also been reported (11–13) and proposed to confer computational advantages (14, 15). How the brain reconciles low-dimensional dynamics with high-dimensional representations in the same neuronal population is an open question.

Here we investigated this question using large-scale recordings of neurons during slow-wave sleep (SWS) to probe the intrinsic population dynamics and examine their relation to activity during awake visual perception. Although previous work has examined replay during SWS (16–18), replay events account for only a small percentage of neuronal activity (19, 20). We aimed instead to use SWS as a window into internally-generated population structure that is uncontaminated by the influence of the ongoing behavior and sensory inputs present during wakefulness (21–23).

## Results

### Neural activity in slow-wave sleep reveals low-dimensional internally-generated population structure

Internally-generated population structure is mostly preserved between sleep and wakefulness (8, 21, 22, 24), but we reasoned that SWS may be closer to a true “ground state” of the brain and could provide better estimates of this population structure. We tested this by first investigating if the population structure observed in sleep is preserved in wakefulness. For this purpose, we performed recordings with long periods of SWS and awake states. We used cross-validated Principal Component Analysis (cvPCA) to estimate the population structure in the data and found that SWS population structure is mostly preserved in awake states (Fig. 1A), with high-variance SWS dimensions accounting for most of the variance in awake population activity (Fig. S1, 98.4 ± 0.7% of awake variance; mean ± s.e.m., n = 130 recordings, multiple brain areas), even after subsampling to account for differences in firing rates (Fig. S2, 89.4 ± 0.6% of awake variance; mean ± s.e.m., n = 130 recordings, multiple brain areas).

**Fig. 1.**
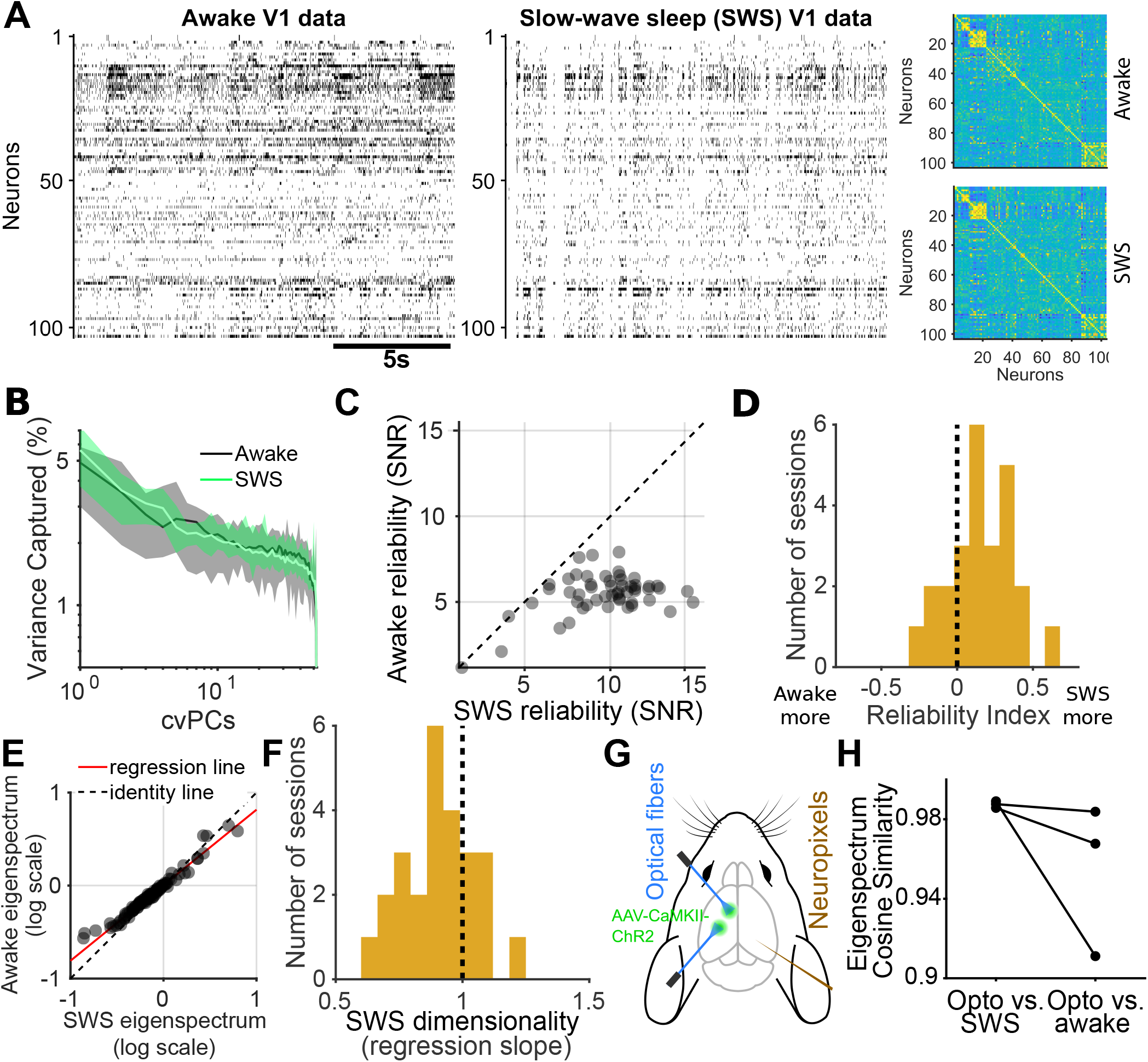
Slow-wave sleep (SWS) enables reliable estimation of internally-generated low-dimensional manifold structure. (**A**) Typical segment of data illustrating that the correlation structure of population activity in V1 is similar during wakefulness and slow-wave sleep (SWS). (**B**) Cross-validated principal components analysis (cvPCA) eigenspectra estimated from five adjacent segments of data show more segment-to-segment variability in wakefulness than in SWS (note wider confidence interval), suggesting SWS provides a more reliable estimate of population structure. (**C**) Individual eigenvalues from panel B are estimated more reliably from SWS than from awake spontaneous activity. (**D**) cvPCA eigenspectra can be estimated more reliably from SWS than wakefulness). (**E**) cvPCA eigenspectra decay faster in SWS than wakefulness (regression slope <1), indicating that SWS is lower-dimensional. (**F**) Linear regression slopes of awake and SWS eigenspectra show that, in general, dimensionality is higher during awake states. (t-test for slope <1) (**G**) We reasoned that if SWS population structure is generated internally, it should be evoked by diffuse, nonspecific optogenetic stimulation. We tested this by stimulating in S1 and M1 while recording in the contralateral hemisphere. (**H**) The eigenspectrum of optostimulation-evoked activity was more similar to SWS than wakefulness, indicating similar population structure

Since SWS is a more homogenous state than wakefulness, we next tested whether SWS activity would provide a more reliable estimate of the intrinsic population structure. To accomplish this, we made independent estimates of the population activity eigenspectrum from short segments of the data (16.83 ± 0.82 min., mean ± s.e.m., n = 141 recordings, multiple brain areas), finding that SWS eigenspectrum estimates showed lower segment-to-segment variability (Fig. 1B-C). Using an index of reliability to quantify this variability (see methods), we found that SWS provides more reliable estimates of population structure than awake spontaneous activity (Fig.1D, reliability index 0.15 ± 0.04, mean ± s.e.m., n = 25 recordings) in both our V1 recordings and in publicly available datasets (Fig. S10) (21, 25–29). We reasoned further that if SWS is closer to a ground state, then activity should be lower-dimensional than in wakefulness. To test this, we compared the rates of eigenspectrum decay (11), finding that SWS activity was lower-dimensional in V1 (Fig. 1E-F, regression slope 0.91 ± 0.03, mean ± s.e.m., n = 25 recordings) and all brain regions tested (Fig. S10). Because REM sleep was higher-dimensional than SWS (Fig. S9), we focused our subsequent analyses on SWS.

Finally, we asked whether the population structure we observed during SWS was truly generated by internal dynamics as opposed to driven by structured inputs such as specific motor commands whose execution is blocked during SWS. To test this, we used Neuropixels probes to record from multiple sites in one hemisphere during wakefulness, SWS, and awake optogenetic stimulation of contralateral M1 or S1 (Fig. 1G). Our rationale was that if SWS population structure is internally-generated, then diffuse, unstructured stimulation during wakefulness should evoke SWS-like activity patterns. In support of our hypothesis, we found that awake optostimulation of the contralateral hemisphere evoked multidimensional dynamics in V1, and the resulting eigenspectrum was more similar to SWS than wakefulness (Fig. 1H, S3, cosine similarity: opto vs. SWS 0.978 ± 0.005, opto vs. awake 0.865 ± 0.027, mean ± s.e.m., n = 8 recordings of multiple brain regions).

### Off-manifold dimensions are activated in wakefulness

For our subsequent analyses, we operationally defined three subspaces containing cvPCs that account for more (“on-manifold”), equal (“non-manifold”), or less (“off-manifold”) variance than chance during SWS (Fig. 2A). Although the high-variance cvPCs do not define a manifold per se, we use the term “on-manifold” to indicate a subspace that likely contains an internally-generated manifold (5, 6, 22, 23). The off-manifold subspace is also noteworthy: these dimensions are preserved in cross-validation and therefore are not merely noise. Instead, they represent patterns of neuronal activity that are reproducibly *less likely than chance* to occur during SWS.

**Fig. 2.**
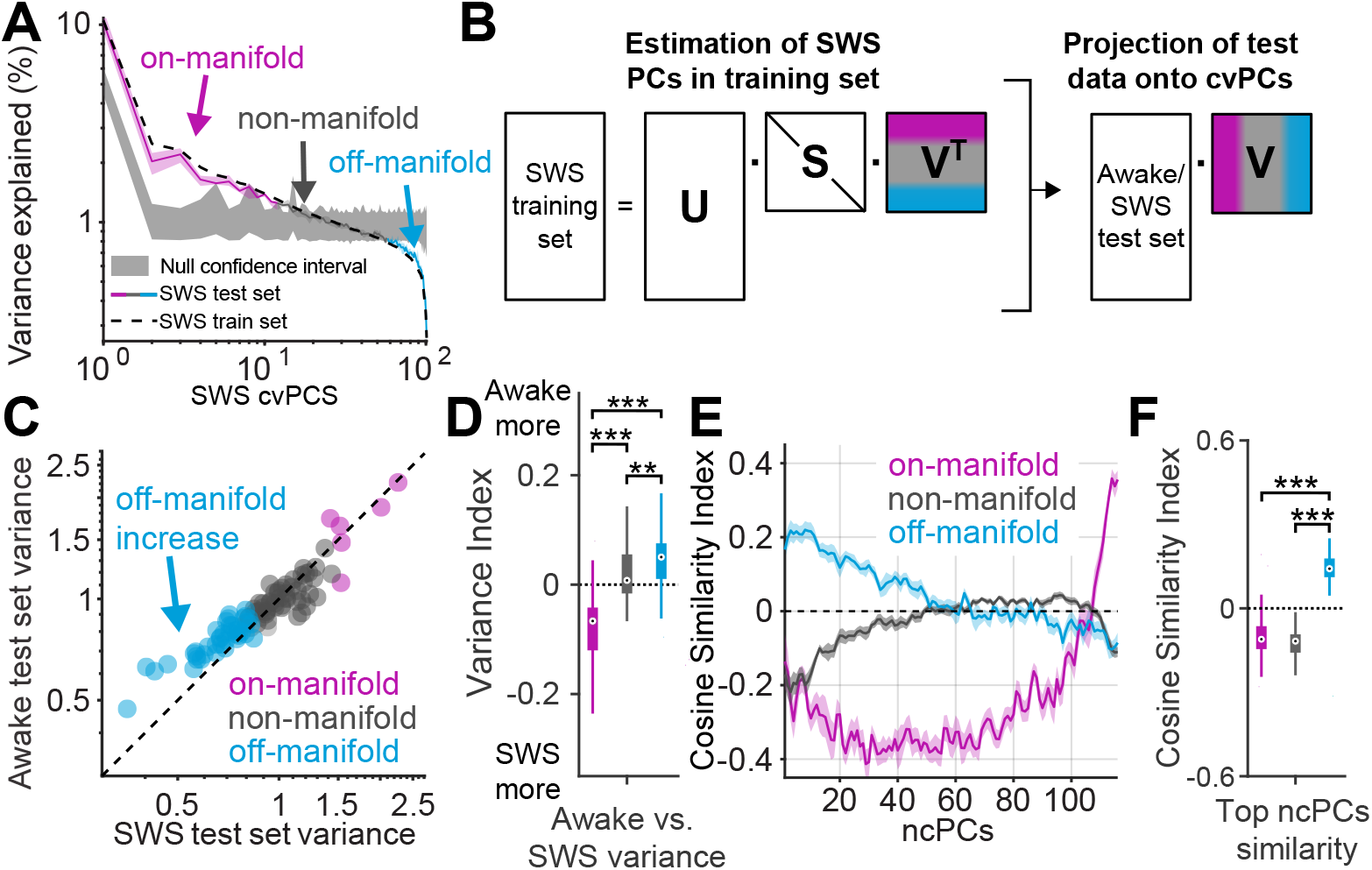
Wakefulness exhibits more off-manifold activity than slow-wave sleep. (**A**) Using SWS cvPCA, we operationally defined three subspaces: on-manifold (magenta), non-manifold (gray), and off-manifold (cyan), which during SWS account for more, equal, or less variance than chance, respectively. Example shows a single session in mouse V1. (**B**) To measure the amount of variance in each subspace, we projected SWS and awake test sets onto cvPCs from SWS training set. (**C**) Awake test set activity shows preferential activation of off-manifold dimensions relative to the SWS test set. (**D**) This result was observed across several V1 sessions analyzed. (**E**) Normalized contrastive PCA (ncPCA, see methods) finds the dimensions that show the largest relative increase in variance between wakefulness and SWS. These dimensions are overrepresented in the off-manifold subspace. (**F**) The top 10% of ncPCs are more aligned with the off-manifold subspace than the on-/non-manifold subspaces [repeated measures ANOVA, multiple comparisons **p<0.01, ***p<0.001].

To examine the similarities and differences between awake and SWS activity, we projected the awake data onto the SWS cvPCs (Fig. 2B) and determined the amount of variance captured by each subspace. We found that awake population structure was similar to SWS, but with increased activity in the off-manifold subspace (Fig. 2C). This increase was observed in several recording sessions in V1 (Fig. 2D, variance index: on-manifold −0.07 ±0.01, non-manifold 0.02 ±0.01, off-manifold 0.04 ± 0.01, mean ± s.e.m., n = 25 recordings), and it was not observed in the opposite direction (SWS projected onto awake cvPCs, Fig. S4), suggesting it is not a trivial consequence of PCA minimizing variance in the off-manifold subspace. To guard further against this possibility, we developed a novel type of PCA, normalized contrastive PCA (ncPCA, see methods), which finds the dimensions exhibiting the largest increase in variance in the awake state. Unlike standard contrastive PCA (30), ncPCA is normalized to remove the bias toward high-variance dimensions. Using ncPCA, we found that the dimensions most overrepresented in wakefulness are preferentially off-manifold (Fig. 2E-F, cosine similarity index: on-manifold −0.096 ± 0.02, non-manifold −0.13 ± 0.01, off-manifold 0.13 ± 0.02, mean ± s.e.m., n = 25 recordings). Together, these results suggest that awake activity is composed of the low-dimensional dynamics observed in SWS with off-manifold activity added. We found similar results in multiple other brain regions (Fig. S10), suggesting this may be a general feature of neural population activity.

### Movements and visual stimuli are both encoded on-manifold, but only stimuli are encoded off-manifold

Since natural scenes have high-dimensional representations in V1 (11), we next investigated how movements and natural visual scenes are encoded in the on/non/off-manifold subspaces (Fig.3A). For this, we projected awake activity into each of the three subspaces and tested their ability to predict movements and stimulus identity compared to random subspaces of equal dimension. In our chronic V1 recordings, we found that running speed is encoded preferentially in the on-manifold subspace (Fig.3B on-manifold 0.18 ± 0.05, non-manifold −0.02 ± 0.01, off-manifold −0.02± 0.02, mean ± s.e.m., n = 6), suggesting that it drives internally-generated dynamics. We also confirmed this result in publicly available datasets collected from several different brain areas (Fig. S10). Further, we found that face motion, a multidimensional movement, is also encoded in the on-manifold subspace in V1 (Fig. 3B on-manifold 0.08 ± 0.02, non-manifold −0.08 ± 0.009, off-manifold −0.1± 0.03, mean ± s.e.m., n = 6 recordings) and several other brain regions (Fig. S10). It has previously been suggested that distributed movement representations are created by efference copy and sensory reafference (2, 31). Although these inputs undoubtedly drive activity, our results indicate that the population structure of movement-related activity is dictated by internal dynamics that are preserved during SWS.

**Fig. 3.**
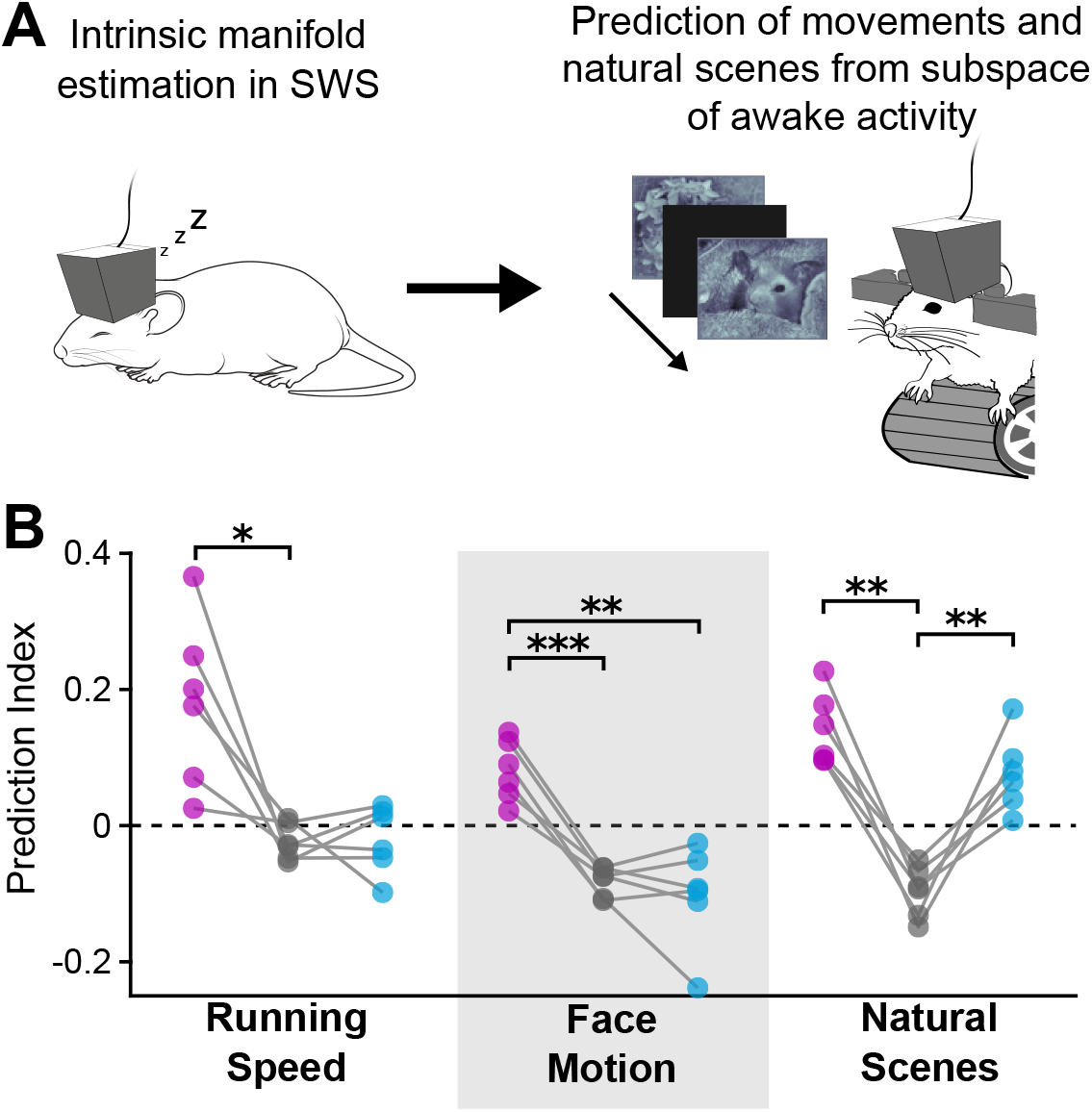
Natural scenes, but not movements, are encoded in the off-manifold subspace. (**A**) We recorded during both SWS and wakefulness in freely-moving mice that were subsequently head-fixed during visual stimulus presentation. We then decoded spontaneous movements and visual stimuli from awake activity projected into the on/non/off-manifold subspaces. (**B**) In V1 recordings, we found that running speed and face motion were encoded on-manifold. However, natural scenes were encoded in both the on- and off-manifold subspaces. The prediction index quantifies prediction performance relative to a random subspace of the same dimensionality. Prediction index: running speed on-manifold 0.20 ± 0.05, non-manifold −0.02 ± 0.01, off-manifold −0.02 ±0.01; face motion: on-manifold 0.06 ± 0.02, non-manifold −0.1 ± 0.008, off-manifold −0.09± 0.03; natural scenes: on-manifold 0.15 ± 0.03, non-manifold −0.1 ± 0.01, off-manifold 0.07 ± 0.02, mean ± s.e.m., n = 6 [repeated measures ANOVA, multiple comparisons *p<0.05, **p<0.01, ***p<0.001].

Unexpectedly, we found that stimulus identity in V1 is encoded in both the on- and off-manifold subspaces, but not the non-manifold subspace (Fig. 3B, S6, S7 on-manifold 0.14 ± 0.02, non-manifold −0.1 ± 0.01, off-manifold 0.08 ± 0.02, mean ± s.e.m., n = 6 recordings). The on-manifold component is consistent with previous studies showing that stimulus-evoked activity in V1 is more similar to spontaneous activity than chance (8, 32–34). However, our results indicate that natural scenes are also encoded by off-manifold patterns of activity that are *less likely than chance* to occur during SWS. Moreover, decoding performance was improved further by using combined on- and off-manifold activity (Fig. S8), indicating that these subspaces encode non-redundant information.

### Off-manifold activity is composed of “chorister” neurons firing decoupled from the rest of the population

We next investigated which neuronal activity patterns occupy the off-manifold subspace. Since natural visual scenes are known to be encoded by sparse activity in V1 (35–37), we first quantified the population sparsity of each SWS cvPC by calculating the Gini coefficient (38). Our analysis revealed that off-manifold cvPCs have higher population sparsity than on- and non-manifold ones (Fig. 4A-B, Gini coefficient: on-manifold 0.45 ± 0.005, n = 82 cvPCs, non-manifold 0.49 ± 0.01, n = 501 cvPCs, off-manifold 0.55 ± 0.003, n = 277 cvPCs, mean ± s.e.m.), meaning that fewer neurons participate in each dimension, the same results was observed in other brain regions (Fig. S10). It has been reported that cortical neurons exhibit a broad distribution of population coupling strengths ranging from “soloists,” which fire decoupled from the rest of the population (i.e. sparse), to “choristers,” whose activity is strongly coupled to the population (i.e. dense) (9) and modulated by movement (Fig. S5). We therefore asked whether the sparse activity patterns we observed in off-manifold dimensions were due to preferential participation by soloist neurons. We found the opposite result: the off-manifold subspace was more likely to contain choristers, and the non-manifold subspace was more likely to contain soloists (Fig. 4C, Pearson correlation: non-manifold r = −0.728 ± 0.015, off-manifold r = 0.729 ± 0.007, mean ± s.e.m., n = 6 recordings). These results were also replicated in other brain regions (Fig. S10), indicating is not V1 specific). The explanation for this seemingly contradictory result is that the off-manifold subspace contains population-sparse activity in neurons with low *average* population sparsity; in other words, it is when chorister neurons fire decoupled from the population (Fig. 4D). Because this is statistically unlikely to occur, it is observed at less-than-chance levels during SWS. In contrast, non-manifold activity occurs at chance levels because soloist neurons usually fire with high population sparsity.

**Fig. 4.**
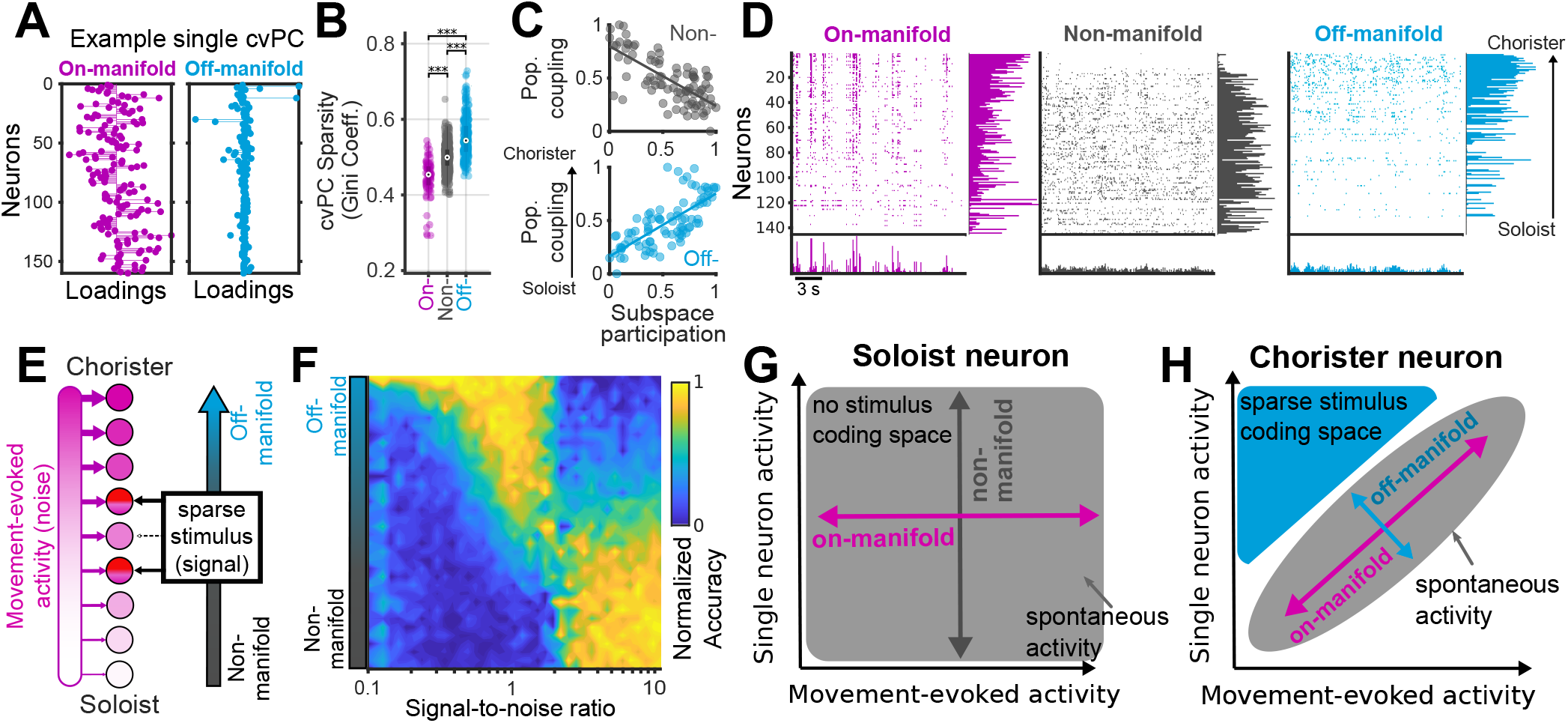
Off-manifold activity consists of population-sparse activity in “chorister” neurons, which usually fire with low population sparsity. (**A**) Representative cvPCs illustrating that fewer neurons participate in a typical off-manifold cvPC (right) than in an on-manifold cvPC (left), indicating higher population sparsity. (**B**) Off-manifold cvPCs have higher population sparsity than on- or non-manifold cvPCs (Gini coefficient, one-way ANOVA, p<0.001). (**C**) Population coupling is a metric of single neuron modulation by overall neural activity. (top) “Soloist” neurons, which have low population coupling and therefore high sparsity, participate more in non-manifold activity. (bottom) “Chorister” neurons, which have high population coupling and low sparsity, participate more in off-manifold activity. Thus, off-manifold activity has high population sparsity but occurs mostly in neurons with low average population sparsity (t-test p<0.001). (**D**) Off-manifold activity is composed of chorister neurons firing sparsely, decoupled from the rest of the population. This panel shows a single segment of V1 data split into its projection into the three subspaces. Rows are ordered by population coupling with choristers at the top and soloists at the bottom. (left) Pseudo-raster plot of on-manifold activity. Note that most activity is near the top of the plot, indicating that choristers show the greatest on-manifold participation, and the bottom histogram has high peaks indicating low population sparsity. (center) Non-manifold activity is mostly in soloist neurons, and the bottom histogram is flat indicating high population sparsity. (right) Off-manifold activity is mostly in chorister neurons, but the population sparsity is high. (**E**) To understand possible advantages of off-manifold coding, we made a model of Poisson-spiking neurons that receive varying amounts of movement-evoked correlated “noise.” Neurons with strong noise inputs fired as choristers (top), and weak noise inputs fired as soloists (bottom). Visual stimuli were encoded by patterns of sparse input to a subset of neurons that was varied from soloists to choristers, thus controlling how much stimulus-evoked activity was in the non- and off-manifold subspaces (Fig. S13). (**F**) For low signal-to-noise ratios, stimulus decoding performance was better when the stimulus was encoded off-manifold in choristers. This indicates, counterintuitively, that it is easier to separate signal and noise when they are encoded in the same neurons rather than different ones. (**G**) An intuitive explanation for this result is that soloist neurons are decoupled from the spontaneous activity manifold, so movement-evoked activity enters the non-manifold space at random, and there is no dedicated space in which to encode the stimulus. (**H**) For chorister neurons, activity is coupled to the manifold, which creates a protected off-manifold subspace (cyan) that movement-evoked activity is unlikely to enter. Sparse activity in chorister neurons accesses this off-manifold coding space, which encodes high-dimensional stimuli separate from movement-evoked activity.

### Low-dimensional dynamics create an off-manifold coding space accessed by sparse activity

These findings suggest that not all sparse activity in V1 is equal: sparse firing carries more information about stimulus identity when it occurs in chorister neurons, which are unlikely to fire sparsely. This implies that visual stimuli are encoded in the neurons with the strongest modulation by movement, which raises the problem that movement-related activity could interfere with visual processing. To gain insight into this, we built a model with Poisson-spiking neurons that receive a dense movement-evoked “noise” input with a distribution of input strengths ranging from low (soloists) to high (choristers) (Fig. 4E). A set of neurons was chosen to encode visual stimuli (“signal”), with each stimulus activating a sparse subset of those neurons. We then systematically varied which set of neurons received visual inputs, ranging from soloists to choristers, enabling us to adjust how much stimulus-related variance was in the non- and off-manifold subspaces (Fig. S13). We found that when the signal-to-noise ratio is high, non-manifold coding by soloists yielded optimal stimulus decoding performance (Fig. 4F). However, when the signal-to-noise ratio is lower (comparable to levels observed in our data: SNR = 1.13 ± 0.03, mean ± s.e.m., n = 6), off-manifold coding by choristers becomes optimal; in other words, encoding movements and stimuli in the same neurons makes it easier to keep them separate. An explanation for this counterintuitive result is that for soloist neurons, spontaneous activity travels throughout the whole state space (Fig. 4G), and stimulus representations overlap with movement-related activity. For chorister neurons, spontaneous activity is constrained to a low-dimensional manifold, opening an unused region of neural state space that sparse activity can access to encode visual stimuli without interference from movement-related activity (Fig. 4G).

## Discussion

This work aims to reconcile several disparate findings in the literature: brain-wide representations of movement (1, 2), low-dimensional dynamics (5–7, 9, 10), and high-dimensional representations of sensory stimuli (11). The population structure of movement-evoked activity has generally been assumed to be generated by motor efference copy or sensory reafference (2, 31). However, we found that this structure is intact during sleep and evoked by diffuse optogenetic stimulation, suggesting that efference copy and sensory reafference merely drive internally-generated dynamics. Our finding that visual scenes are encoded partly in the on-manifold subspace suggests that structured sensory inputs also activate internally-generated dynamics, consistent with previous studies showing that evoked and spontaneous activity in V1 are more similar than expected by chance (32–34). However, we unexpectedly found that stimulus identity is also encoded in the off-manifold subspace, which contains activity patterns that occur at less-than-chance levels. This suggests that structured inputs can evoke high-dimensional population activity patterns that are not generated spontaneously, providing one possible explanation for how high-dimensional representations can be compatible with low-dimensional dynamics and movement-related activity. Whether these off-manifold patterns passively reflect the structure of their inputs or are stored in the local network and triggered by structured inputs will be an interesting question for future studies. Our finding that off-manifold activity is composed of population-sparse activity in chorister neurons, which are the least likely to fire sparsely, reveals an unexpected link between dimensionality and sparse coding. Sparse representations have been reported in numerous brain regions of different species (35, 39–41), where they have been proposed as a mechanism to reduce energy consumption (42). Our results suggest that another important function of sparse coding may be to access the off-manifold subspace. Conversely, an underappreciated function of the low-dimensional dynamics found in many brain regions may be to reserve an off-manifold coding space for high-dimensional representations. Our finding that wakefulness is associated with increased off-manifold activity in many brain regions suggests this may be a general principle that holds outside of V1. This study has several important limitations, including the fact that we did not attempt to use nonlinear dimensionality reduction methods to determine the intrinsic dimension or structure of the underlying population activity manifold. Additionally, we did not attempt to determine which specific stimulus features are coded in the on- and off-manifold subspaces. Future studies addressing these questions could shed light on the possible roles of off-manifold activity in encoding higher-order stimulus features or unexpected deviations from movement-evoked changes in sensory inputs. Our findings highlight the benefits of using sleep as a tool to study awake brain function, even in primary sensory areas such as V1. Many previous studies have compared evoked activity to awake spontaneous activity (9, 10, 32, 34, 43, 44), but our results suggest that SWS provides a more reliable estimate of internally-generated low-dimensional population structure. We encourage others to incorporate sleep recordings into their experiments as a relatively low-effort means to gain insight into the interactions between internally-generated dynamics and representations of sensory and cognitive variables.

## ACKNOWLEDGMENTS

We wish to thank Ruben Coen-Cagli, Adam Kohn, Dan Levenstein, Michael Long, and Jacob Ratliff for helpful comments on the manuscript. We also wish to thank György Buzsáki, in whose lab the published datasets used in this study were collected. This work was supported by funds from the National Institute on Drug Abuse (DA051608 and DA036657), the Leon Levy Foundation, the Brain and Behavior Research Foundation, and Montefiore/Einstein startup funds.

## AUTHOR CONTRIBUTIONS

L.S. and A.P. conceived the project. E.F.O., S.K., R.B.B., and L.S. designed the experiments. E.F.O. collected all freely-moving electrophysiological data, and S.K. collected all head-fixed Neuropixels data. E.F.O., S.K., T.S.Q., and L.S. performed data analysis, and S.K. and E.F.O. performed computational modeling. L.S. supervised all aspects of the project. E.F.O., S.K., and L.S. wrote the manuscript with contributions from all authors.

## Methods

All experimental procedures were conducted in accordance with the Institutional Animal Care and Use Committee of Albert Einstein College of Medicine.

### Animals and surgery

Experiments were conducted using C57BL/6J x FVB F1 hybrid mice (45). For the visual stimulation task, we chronically implanted a 64-site silicon probe (NeuroNexus) in the primary visual cortex (V1) (AP: 3.4 ML: 2.6 DV: 0.6 from brain surface) under isoflurane anesthesia, as described previously (46). Wires for reference and electrical ground were implanted above the cerebellum, and a copper mesh hat was built to shield the probes. Probes were mounted on microdrives that were used to move the probe farther into V1 for maximizing unit yield, though never beyond DV of 1.0 mm. Animals were housed individually after implantation and allowed to recover for at least one week before experiments. Recordings were performed using the Intan system at 30 kHz. Offline automatic spike sorting was performed using Kilosort 2.0 (1, 47), and all parameters used for sorting are presented in the [Kilosort 2.0 wrapper](https://github.com/eliezyer/KS2wrapper) repository in StandardConfig_KS2Wrapper.m. Manual adjustment of Kilosort outputs was performed using [Phy](https://github.com/cortex-lab/phy). Isolated single units were assigned to putative principal neurons or fast-spiking interneurons based on through-to-peak time of waveforms, a criteria previously used in V1 (28).

We also used data that is publicly available on the [Buzsáki laboratory website](https://buzsakilab.nyumc.org/datasets/). We used the Matlab toolbox [Buzcode](https://github.com/buzsakilab/buzcode) to extract the data used for this manuscript. The methodology for collection of this data was described elsewhere (21, 25–29). From all datasets, we selected only recording sessions where there were location/speed tracking and slow-wave sleep (SWS) states. A table with a brief summary of the dataset used is presented in table 1.

**Table 1.**
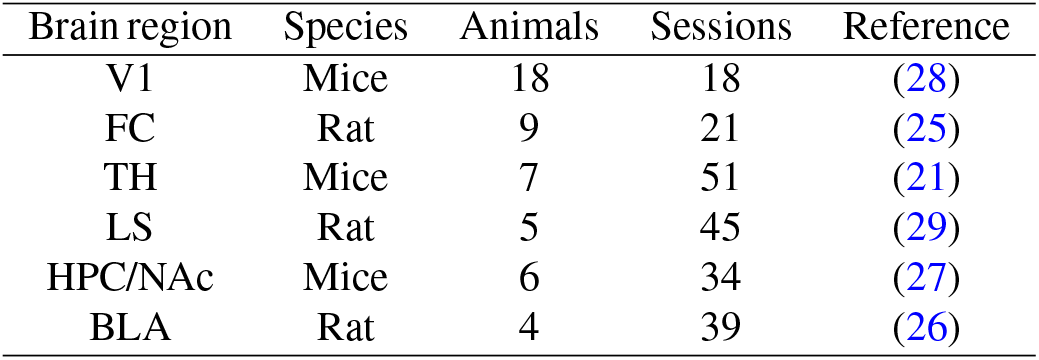
Summary of datasets used for analysis. V1: primary visual cortex, FC: frontal cortex, TH: thalamus, LS: Lateral Septum, HPC: Hippocampus, NAc: Nucleus Accumbens, BLA: Basolateral amygdala

**Table 2.**
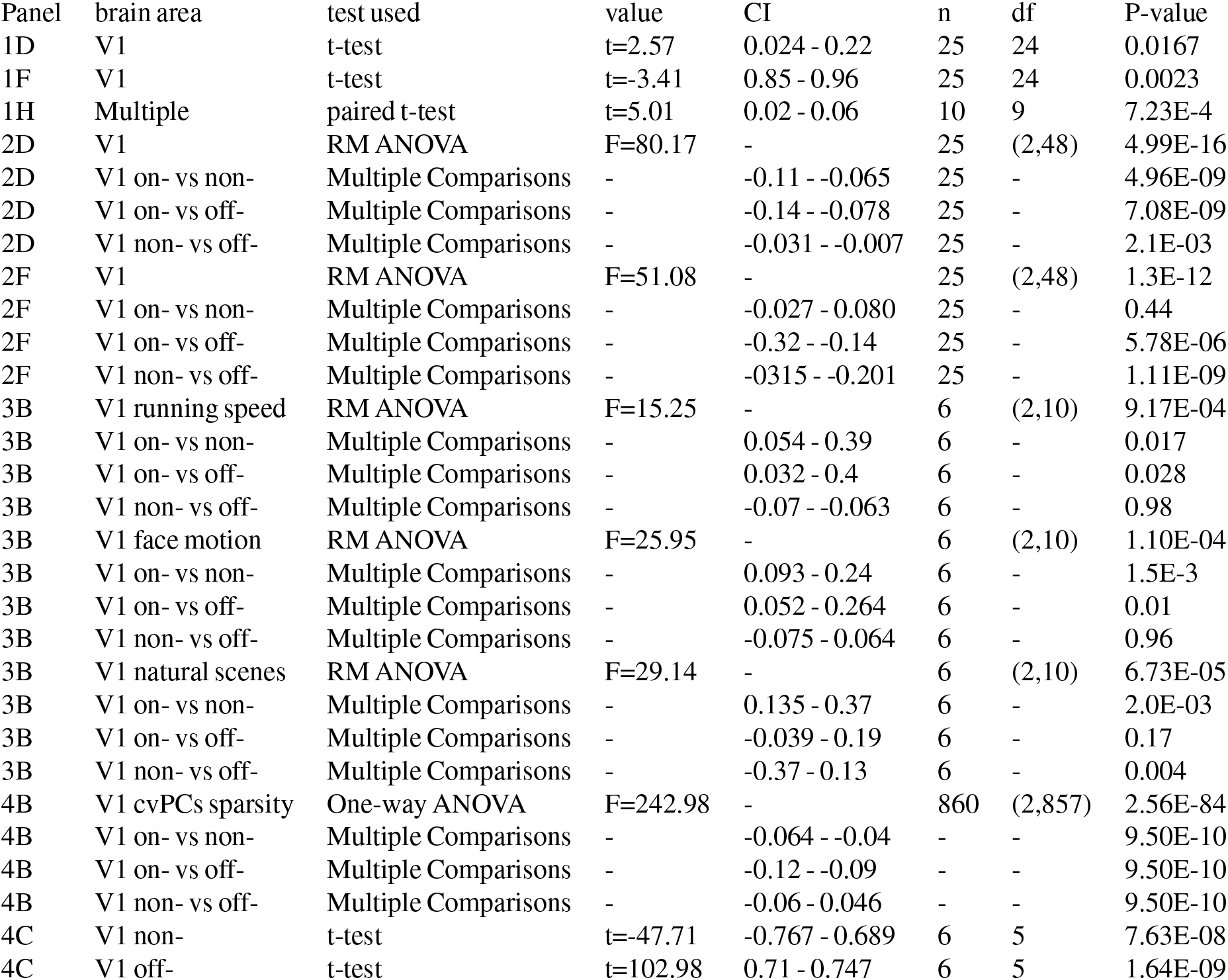
Summary of statistics for the main figures.

| Panel | brain area | test used | value | CI | n | df | P-value |
| --- | --- | --- | --- | --- | --- | --- | --- |
| 1D | V1 | t-test | t=2.57 | 0.024 - 0.22 | 25 | 24 | 0.0167 |
| 1F | V1 | t-test | t=-3.41 | 0.85 - 0.96 | 25 | 24 | 0.0023 |
| 1H | Multiple | paired t-test | t=5.01 | 0.02 - 0.06 | 10 | 9 | 7.23E-4 |
| 2D | V1 | RM ANOVA | F=80.17 | - | 25 | (2,48) | 4.99E-16 |
| 2D | V1 on- vs non- | Multiple Comparisons | - | -0.11 - -0.065 | 25 | - | 4.96E-09 |
| 2D | V1 on- vs off- | Multiple Comparisons | - | -0.14 - -0.078 | 25 | - | 7.08E-09 |
| 2D | V1 non- vs off- | Multiple Comparisons | - | -0.031 - -0.007 | 25 | - | 2.1E-03 |
| 2F | V1 | RM ANOVA | F=51.08 | - | 25 | (2,48) | 1.3E-12 |
| 2F | V1 on- vs non- | Multiple Comparisons | - | -0.027 - 0.080 | 25 | - | 0.44 |
| 2F | V1 on- vs off- | Multiple Comparisons | - | -0.32 - -0.14 | 25 | - | 5.78E-06 |
| 2F | V1 non- vs off- | Multiple Comparisons | - | -0.315 - -0.201 | 25 | - | 1.11E-09 |
| 3B | V1 running speed | RM ANOVA | F=15.25 | - | 6 | (2,10) | 9.17E-04 |
| 3B | V1 on- vs non- | Multiple Comparisons | - | 0.054 - 0.39 | 6 | - | 0.017 |
| 3B | V1 on- vs off- | Multiple Comparisons | - | 0.032 - 0.4 | 6 | - | 0.028 |
| 3B | V1 non- vs off- | Multiple Comparisons | - | -0.07 - -0.063 | 6 | - | 0.98 |
| 3B | V1 face motion | RM ANOVA | F=25.95 | - | 6 | (2,10) | 1.10E-04 |
| 3B | V1 on- vs non- | Multiple Comparisons | - | 0.093 - 0.24 | 6 | - | 1.5E-3 |
| 3B | V1 on- vs off- | Multiple Comparisons | - | 0.052 - 0.264 | 6 | - | 0.01 |
| 3B | V1 non- vs off- | Multiple Comparisons | - | -0.075 - 0.064 | 6 | - | 0.96 |
| 3B | V1 natural scenes | RM ANOVA | F=29.14 | - | 6 | (2,10) | 6.73E-05 |
| 3B | V1 on- vs non- | Multiple Comparisons | - | 0.135 - 0.37 | 6 | - | 2.0E-03 |
| 3B | V1 on- vs off- | Multiple Comparisons | - | -0.039 - 0.19 | 6 | - | 0.17 |
| 3B | V1 non- vs off- | Multiple Comparisons | - | -0.37 - 0.13 | 6 | - | 0.004 |
| 4B | V1 cvPCs sparsity | One-way ANOVA | F=242.98 | - | 860 | (2,857) | 2.56E-84 |
| 4B | V1 on- vs non- | Multiple Comparisons | - | -0.064 - -0.04 | - | - | 9.50E-10 |
| 4B | V1 on- vs off- | Multiple Comparisons | - | -0.12 - -0.09 | - | - | 9.50E-10 |
| 4B | V1 non- vs off- | Multiple Comparisons | - | -0.06 - 0.046 | - | - | 9.50E-10 |
| 4C | V1 non- | t-test | t=-47.71 | -0.767 - 0.689 | 6 | 5 | 7.63E-08 |
| 4C | V1 off- | t-test | t=102.98 | 0.71 - 0.747 | 6 | 5 | 1.64E-09 |

### Viral vectors

We used an adeno-associated viral vector (AAV) with a CaMKII promoter to express channelrhodopsin-2 in pyramidal cells in primary sensory and motor cortex. The recombinant AAV vector was pseudotyped with AAV5 capsid protein and packaged by Addgene.

### Visual stimuli

Natural scene stimuli were the same as used in (1). In brief, 32 images were selected from ethological relevant categories, such as animals and plants. The images chosen were less than 50% uniform background, and with a balance of low and high spatial frequencies. We performed 60 repetitions with 0.5 s duration and an inter-stimulus interval varying from 0.7 to 1 seconds. Visual stimuli were presented on a screen facing the eye contralateral to the side being recorded.

### Neuropixels data acquistion

Neuropixels electrodes were used to record extracellularly from neurons in multiple brain areas including V1, hippocampus, thalamus, striatum, and prefrontal cortex in head-fixed mice. On the day of recording, two small craniotomies were made with a dental drill. After recovery, animals were head-fixed on a custom made treadmill. Two neuropixels probes are inserted in each animal’s right hemisphere (anterior-posterior (AP): −3.2 mm, medial-lateral (ML): 2.7 mm, depth(D): 3.6 mm, angle: horizontal 60 degrees, medial 15 degrees; AP: −0.2 mm, ML: 3.2 mm, D: 5.6 mm, angle: horizontal 60 degrees, medial 55 degrees; AP: 0.9 mm, ML: 1.1 mm, D: 3.8 mm, angle: horizontal 60 degrees, medial 55 degrees). Each probe was mounted on a rod held by a micromanipulator (uMP-4, Sensapex Inc.) and advanced slowly (~ 1*μ*m/s). Electrodes were allowed to settle for 30 min before starting recording. Recordings were made in external reference mode with LFP gain 250 and AP gain 500 using Open-Ephys software. Wires for reference and electrical ground were connected to the cerebellum.

### Optogenetic stimulation

In each animal’s left hemisphere, the optical fibers (400 μm core diameter, 0.5 NA) were inserted in S1 (anterior-posterior (AP): −1.5 mm, medial-lateral (ML) 1.5 mm, Depth(D): 0.6 mm) and M1 (AP: 1.5 mm, ML 1.7 mm, D: 1.2 mm). Optogenetic stimulation was at 450 nm (Osram PL450B laser diode) with power ranging from 0-6 mW. We use three types of optogenetic stimuli: 25 Hz white noise lasting 1 s, 10 Hz white noise lasting 1 s, and a 10 Hz sinusoid lasting 0.4 s.

### Data analysis and exclusion

We used custom MATLAB custom scripts for analysis and plotting. In our analysis we used the following toolboxes: [Buzcode](https://github.com/buzsakilab/buzcode), [communication subspace](https://github.com/joao-semedo/communication-subspace) (48) for Reduced Rank Regression, and [GPML toolbox](http://gaussianprocess.org/gpml/code/matlab) for Gaussian Process Regression. We used only sessions containing a minimum of 30 neurons and 20 minutes of sleep, unless otherwise stated. When analyzing running speed and face motion prediction, we excluded sessions where the full rank prediction was smaller than a threshold of *R*^2^ = 0.1. In the classification of natural visual scenes we excluded low-quality sessions in which the total prediction accuracy was below 30%.

### Cross-validated PCA

First, we separate the data into training and test sets (*N_tr_* and *N_ts_* respectively). The singular vectors *V_n_* are calculated for the training set, and test set data is projected onto those singular vectors to estimate cross-validated scores *U_n_ = N_ts_V_n_*, with variance calculated from each cvPC score. When estimating the amount of variance, we performed cvPCA in five contiguous folds, i.e. folds that have the same duration but consist of continuous blocks of time rather than randomly selected time points. We set each neuron to contribute equally to the cvPCA by z-scoring them individually. Each state (awake and SWS) is z-scored separately. Since shortening the data in time can change the variance of each neuron, we z-scored the training set separately before PCA, but the test set was only z-scored once with the entire dataset.

To build a null distribution for the eigenspectrum, we shuffled the cell identities of the test set 10,000 times and calculated the variance of the shuffled test set’s projection onto the original singular vectors. We defined the confidence interval as the lower (100 × *p*)% and upper (100 × (1 – *p*))% values of this null distribution, where *p* = 1/*N* and N is the number of units in the data. The confidence interval was used to define where on-/non-/off-manifold subspaces begin and end in the cross-validated eigenspectrum.

### Normalized contrastive PCA

To determine the dimensions that are maximally different between SWS and awake states, we developed a new PCA method to find the contrastive dimensions between the two. In order to control for high variance dimensions dominating the contrastive PCs, we normalize the method by their variance. This method finds the eigenvector *x* that maximizes equation 1.

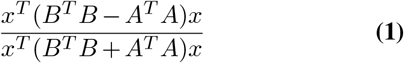

Where *A* and *B* are the neural activity in SWS and awake states.

### Reliability index

The reliability index was calculated based on the coefficient of variation from the cross-validated eigenspectrum. For this purpose, we took segments of awake and SWS data of equal duration and separated the data from each state into five contiguous folds. We performed PCA on one fold and projected each other fold in the testing sets onto the training set PCs. Each fold got a turn as the training set, which allowed us to build a confidence interval out of the PCs. Using the standard deviation (*σ*) and mean (*μ*) we calculated the Reliability index by the ratio 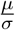 which is equivalent to the inverse of the coefficient of variation 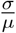.

### Gaussian process regression and running speed prediction

We used gaussian process regression (GPR) to predict running speed from the neural activity. Running speed was log-transformed and z-scored before model training and prediction. The parameters used for GPR were mean function zero, covariance function rational quadratic, and exact inference of posterior probability with gaussian likelihood. All the hyper-parameters were optimized by the log marginal likelihood.

### Face motion extraction and prediction

We record the face of the mouse using a camera with a infrared filter. Infrared LEDs illuminated the face of the mouse. Videos were collected at 30 Hz and aligned with electrophysiology based on digital pulses that triggered frame collection.

To extract the face motion variables we used [Facemap](https://github.com/MouseLand/facemap). For face motion variables used here, we excluded ROIs around the eyes to avoid contamination by the pupil.

Reduced rank regression was used for prediction of face motion components by z-scored neural activity. The initial *β* used for the regression were estimated by ridge regression, as described previously (48).

### Optostimulation prediction and Partial Least Squares regression

To determine the dimensionality of optostimulation-driven population activity we used partial least squares (PLS) regression. This method identifies dimensions in the neural data that maximize the prediction of the optostimulation waveform. Similarly to PCA, PLS regression identifies orthogonal dimensions with decaying variance explained in a successive manner. We performed prediction of optostimulation by using the z-scored neural data projected into the identified PLS dimensions and using GPR to predict the z-scored optostimulation waveform. For GPR we used the same parameters as described in the Gaussian process regression section. This method was done separately for S1 and M1 optostimulation.

To define the dimensionality of optostimulation in the neural data, we picked the number of PLS dimensions that plateaued the cross-validated *R*^2^. For that we picked the lowest number of dimensions that had prediction performance one s.e.m. away from peak prediction.

### Natural visual scenes classification

We made predictions of stimulus identity by fitting a linear multiclass support vector machine (SVM); the SVM model fits a one-vs-one multiple binary classifications to a total of K(K−1)/2 models, where K is the number of visual stimuli identity. The average firing rate of neurons during each stimulation was z-scored and used as a variable for prediction. We projected this data in different dimensions, when stated so, for some of the analysis performed in this study

### R^2^ and accuracy

To estimate performance of regression analysis, we use the equation 2 for *R*^2^.

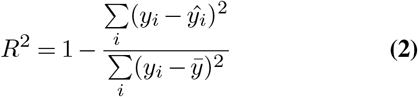

Where *y* is the original variable, *ŷ* is the model predicted and *ӯ* is the mean of the original variable. In this *R*^2^ equation, the model is compared to a constant model (the mean of the original variable). If the prediction is worse than the constant model, the *R*^2^ can be negative, which is comparable to *R*^2^ = 0.

The performance of classification for natural scenes was calculated using the accuracy equation 3.

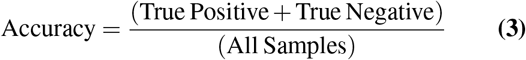

### Prediction index and random projections

There was a different number of dimensions in each subspace (on-/non-/off-manifold), which can influence the prediction quality of the variables we tested. To control for that, we generated 10 random subspaces of equal dimensionality to the on-/non-/off-manifold subspaces. We projected the data into both the original subspace and the random subspaces to make predictions of running speed, face motion, and natural scenes. We used the *R*^2^ from the original subspace and the average *R*^2^ from the subspaces generated to calculate the prediction as indicated in equation 4.

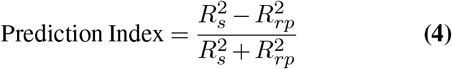

Where 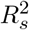 is the *R*^2^ of the subspace and 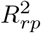 the *R*^2^ random projections of matching dimensionality. A positive prediction index means the prediction was better by the original subspace, and a negative index means that random subspaces generated a better prediction. Because the *R*^2^ cannot be negative for this index calculation, we zeroed every *R*^2^ that was negative (see *R*^2^ and accuracy section).

### Single-neuron population coupling

Population coupling was calculated as described before (9); in brief, we summed the neural activity of N-1 neurons binned and zero-centered, then took the dot product of the left-out neuron’s activity with that of the summed population activity. The resulting number is normalized by the L2-norm of the left-out neuron’s activity. We normalized the population coupling to the range [0,1]. Unlike Okun et al., we did not smooth the data prior to the population coupling calculation.

### Population sparseness

We used the Gini coefficient of the absolute loadings to estimate the population sparseness. This was calculated separately for each cvPC. We first calculated the histogram of loadings using 30 bins, then estimated scores *S_n_* by multiplying the count in each bin by its center value. The histogram of loadings was also used to calculate the discrete density function *f*(*y_i_*), which represents the fraction of the population that has that loading value. Using these variables, we estimated the Gini coefficient by equation 5.

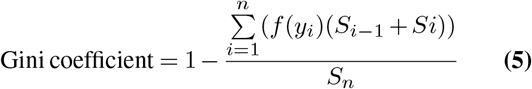

The coefficient varies from 0 to 1, where 0 is perfect equality (i.e. equal participation of all the neurons in a cvPC) and 1 is maximal inequality (only one neuron participates in the cvPC).

### Overall firing rate fluctuation

To control for population firing rate fluctuations in our results, we removed it from the data when applicable. For that we estimated the overall firing rate fluctuation as the change in activity in a [**1**] vector of the same number of rows as the number of neurons. This vector is normalized by its *L*2-norm and used to project the z-scored neural data *N* (organized with time bins in rows and neurons in columns) and estimate the overall firing rate fluctuation in time FR_*t*_. We then removed this from the data by subtracting this projection, as in *N* – FR*t* · [**1**]^**T**^.

### Up and down state detection

To control for up and down transitions in SWS affecting our estimates of population activity, we detected up/down states and restricted the analysis to up states alone. For detecting these states, we used the same methodology as described previously (49). In brief, during periods of SWS, the LFP delta (0.5-8 Hz) and gamma bands (30-600 Hz) are used for thresholding and detection of up states (high gamma power) and down states (delta peak and low gamma power). Down states are then further restricted to periods where spiking activity is below threshold.

### High firing rate spikes subsampled

To control for the possibility that some of our results were biased by differences in neuronal firing rates, we subsampled the spike trains of high firing rate neurons to match the number of spikes in the low firing rate neurons. We sorted the neurons according to their firing rate and subsampled the top half of neurons to match the bottom half. We randomized which top neuron would match which bottom neuron so that the first top neuron did not always match the first bottom neuron.

### Single neuron subspace participation

To determine the participation of a neuron to a specific subspace, we calculated the average absolute loading of that neuron in all the cvPCs that span that specific subspace.

#### Simulation

We built a simple neural network to investigate how the overlap of both movement-evoked activity and visual stimuli influences the decoding of the visual stimulus identity in relation to the signal-to-noise ratio. The network has 100 principal neurons, and each neuron’s firing patterns are Poisson-distributed with uncorrelated noise added at each time step. Neurons were not connected to each other but received inputs encoding both movement “noise” and visual stimulus “signal”, each with their own weight matrix. There are two distinct time epochs: a sleep epoch receiving only the movement-evoked white noise input, and a visual stimulus epoch receiving both the noise input and the visual stimulus signal input. For both epochs we used 100 ms time steps, and the total duration of each epoch is 50 s. During the sleep epoch, the white noise temporal profile was fed into the 60 out of the 100 principal neurons with a weight vector of normal distribution loadings in the range [0,1]. During the visual stimulus epoch, principal neurons also received the same white noise input as during the sleep epoch. In addition, ten different sparse input weight vectors representing ten visual stimuli were fed into the principal neurons with loadings in the range [−1,1]. 30 out of 100 neurons received visual stimuli, each with its own weight vector. 5 out of 30 neurons received all 10 visual stimulus inputs, and different combinations of 5 out of 25 neurons were stimulated with each visual stimulus input. Thus 10 different combinations of neurons are tuned for individual stimuli. In the simulations, two parameters were explored: the signal-to-noise ratio and the signal/noise population overlap. The signal-to-noise ratio is varied by adjusting the amplitude of the white noise stimulus. The signal/noise overlap is defined by selecting 30 out of 100 neurons to receive the visual stimulus, and those neurons vary from having the highest to lowest movement-evoked activity. To test the network’s performance encoding visual stimuli, prediction accuracy was computed using the accuracy formula above.

#### Null hypothesis testing

Information on statistical tests can be found on supplementary table 2. All t-tests were two-tailed, and for multiple pairwise group comparisons we used Tukey-Kramer posthoc test correction.

---

**Fig. S 1.**
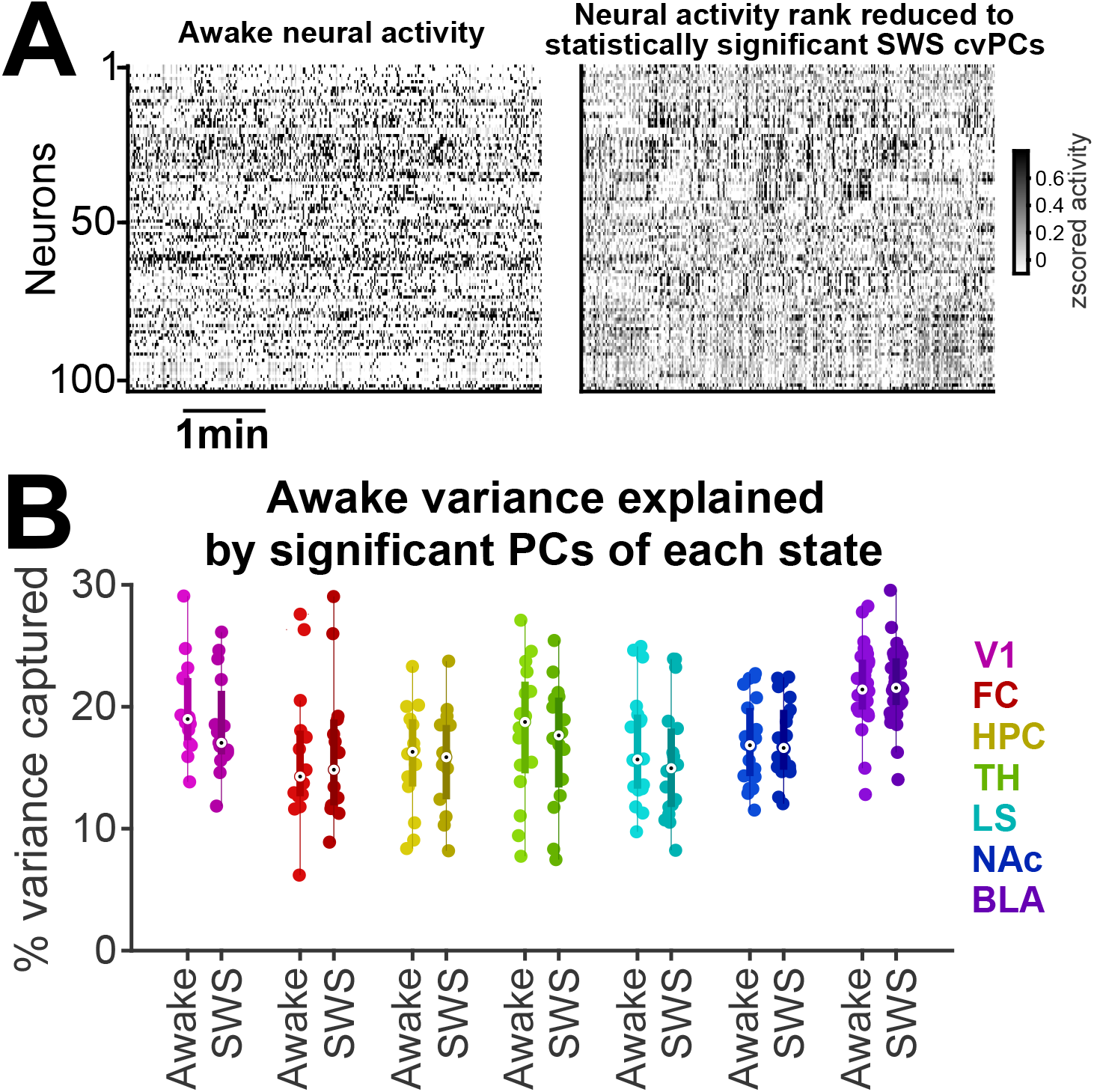
SWS and awake cvPCs explain a similar amount of variance in awake population activity. (**A**) SWS cvPCs capture much of the population structure of awake activity. (**B**) The top k (k = 9.4 ± 0.6, mean ± s.e.m., n = 130 recordings across multiple brain regions) cvPCs of awake and SWS activity (defined as the number of cvPCs with variance above chance in SWS) explain similar amounts of variance in the awake test set data. This suggests that low-dimensional population structure is conserved across SWS and wakefulness.

**Fig. S 2.**
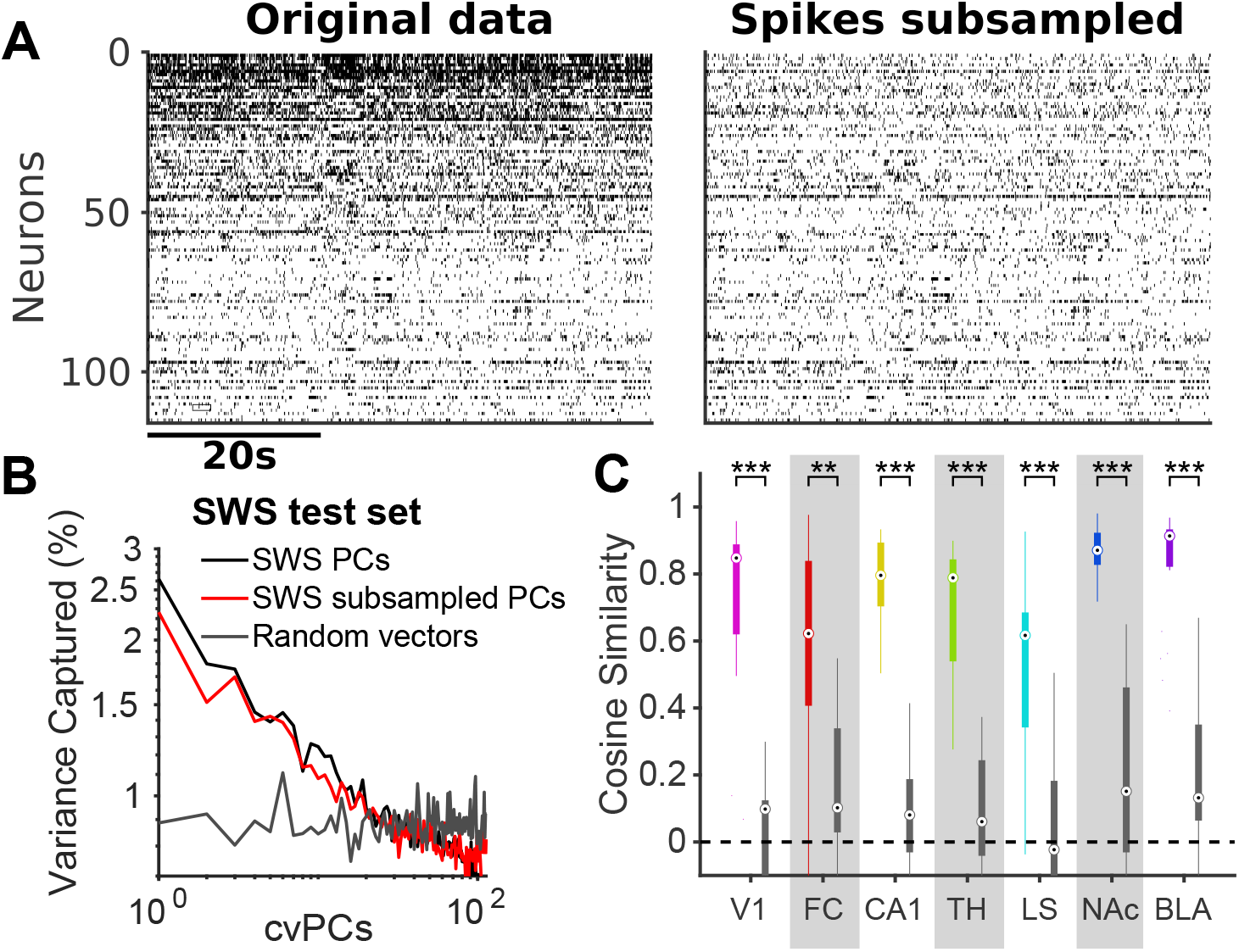
The population activity structure observed during SWS is not explained by high firing rate neurons dominating on-manifold dimensions. (**A**) To control for higher firing rate neurons being overrepresented in on-manifold dimensions, we subsampled the spikes of the top half of high-firing-rate neurons to match the firing rates of the bottom half. (**B**) Example eigenspectrum of original SWS test set projected onto PCs from SWS training set, subsampled SWS training set, and random vectors. Note similarity of subsampled and full eigenspectra. (**C**) The SWS PC eigenspectrum is more similar to the subsampled SWS PC eigenspectrum (colored boxplots) than random vectors (grey boxplots) for all brain areas tested. Eigenspectrum cosine similarity with SWS subsampled: 0.73 ± 0.02, random vectors: 0.12 ± 0.02, mean ± s.e.m., n = 130 recordings. [paired t-test **p<0.01, ***p<0.001]

**Fig. S 3.**
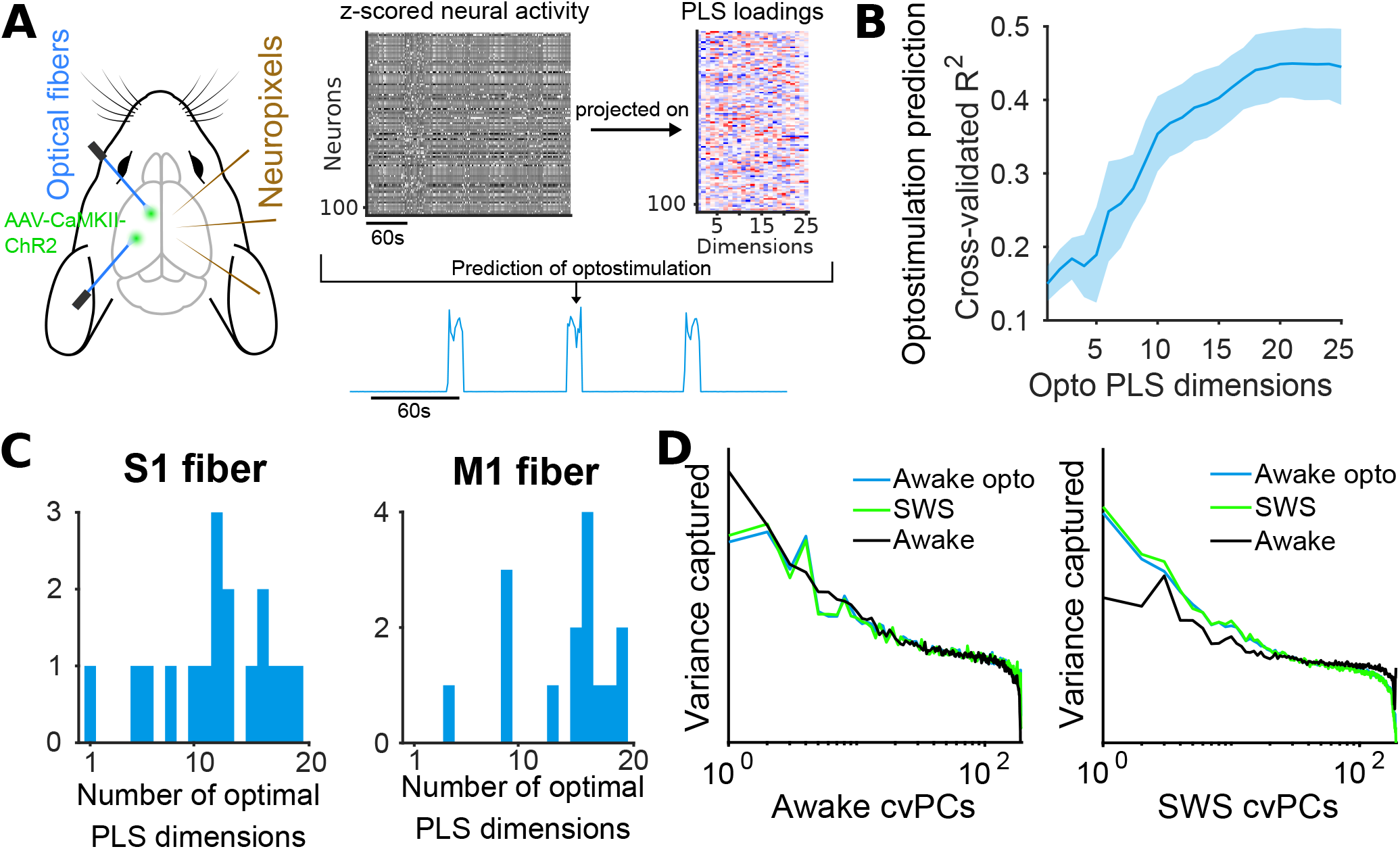
Optogenetic stimulation evokes multidimensional activity patterns similar to those observed during SWS. (**A**) Using partial least squares regression (PLS), we found neuronal activity dimensions that best predicted the white noise waveform of diffuse optogenetic stimulation applied to the contralateral hemisphere. (**B**) Multiple PLS dimensions are required to predict the one-dimensional optogenetic stimulus, suggesting that optostimulation drives internally-generated dynamics. (**C**) S1 and M1 optostimulation generate multidimensional representations in multiple brain areas (S1: 12 ± 1.2, n = 17 brain areas; M1: 14.1 ± 1.2, n = 15 brain areas; mean ± s.e.m.). (**D**) Optostimulation during awake states evokes population activity that is more similar to SWS than awake states. The left eigenspectrum example shows the projection of test set data onto awake cvPCs, and the right eigenspectrum shows the projection onto SWS cvPCs. In both cases, optostimulation is closer to SWS than awake (group statistics shown in Fig. 1H).

**Fig. S 4.**
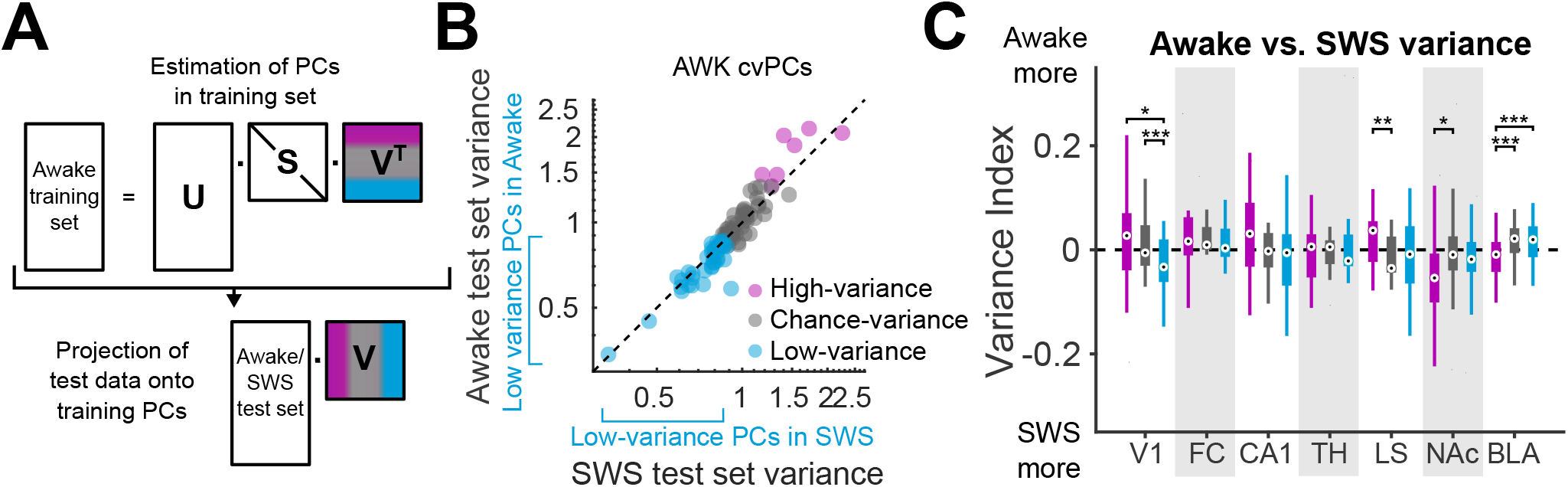
SWS activity does not show increased variance in dimensions that exhibit the lowest variance during wakefulness. (**A**) To test whether our results from Fig. 2 were artifactual, we performed the same analysis in the opposite direction: PCs were estimated from the awake training set, then awake and SWS test sets were projected onto these PCs. We built a null distribution using the awake training set to identify cvPCs that explain more, equal, or less variance than chance. (**B**) The SWS test set projected onto awake cvPCs does not show an increased variance in the low-variance dimensions, as is observed when an awake test set is projected onto SWS cvPCs (Fig. 2C-D). This suggests that the increased off-manifold activity observed during wakefulness is not a trivial consequence of the awake and SWS eigenspectra being different. (**C**) With the sole exception of the BLA, we did not observe any increased variance in the lowest-variance awake PC subspace. Variance index: high-variance −0.002 ± 0.007, chance-variance 0.003 ± 0.005, low-variance 0.01 ± 0.01, mean ± s.e.m., n = 141 recordings). [repeated measures ANOVA, multiple comparisons *p<0.05, **p<0.01, ***p<0.001].

**Fig. S 5.**
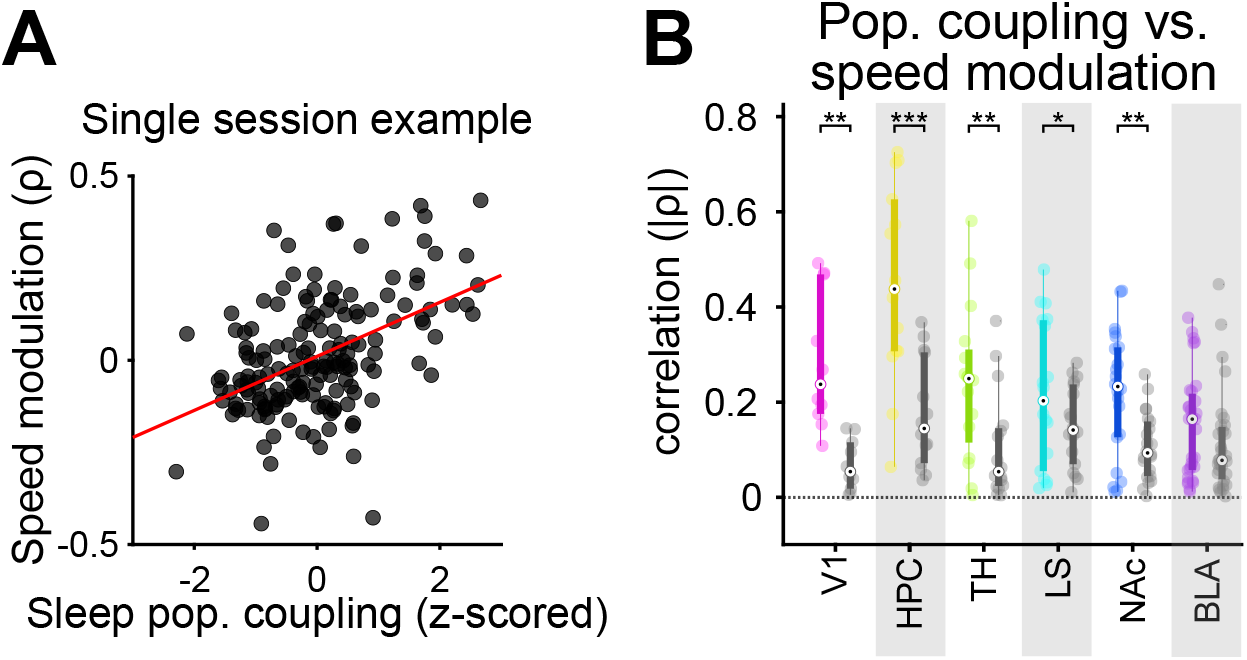
Population coupling during sleep is correlated with running speed speed modulation during wakefulness. (**A**) An example from a single recording session showing that sleep population coupling and awake running speed modulation are correlated. Each dot is a neuron. (**B**) Summary of several brain areas. The absolute value of the correlation coefficient for sleep population coupling and speed modulation for each brain area (colored boxplot) is shown alongside the result from a null distribution where we shift running speed in time (gray boxplot). [paired t-test *p<0.05, **p<0.01, ***p<0.001]

**Fig. S 6.**
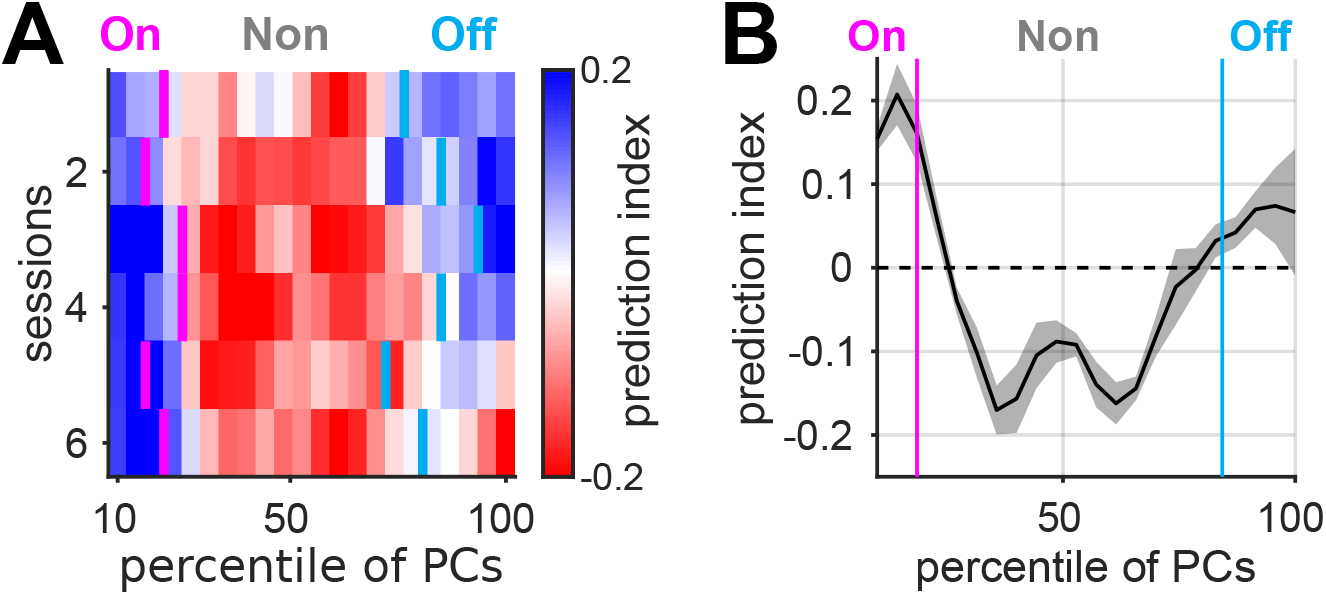
SWS on- and off-manifold boundaries roughly match changes in the encoding of natural scenes in V1. (**A**) On- and off-manifold PCs predict the identity of natural scene stimuli better than non-manifold PCs. (**B**) The averaged prediction index across all sessions shows that changes in decoding performance roughly match the on- and off-manifold boundaries, represented by magenta and cyan lines, respectively.

**Fig. S 7.**
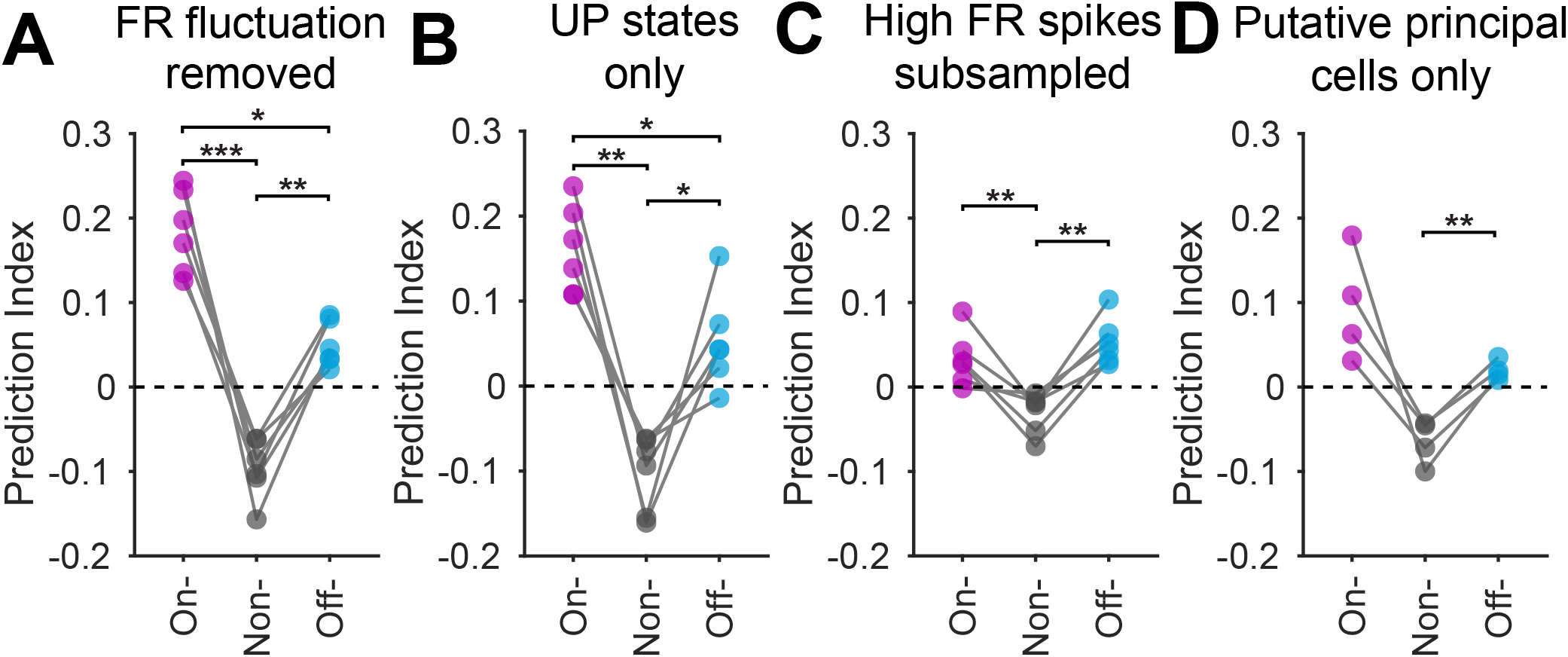
Off-manifold encoding of natural visual scenes is not explained by various nonspecific effects. Off-manifold coding remains intact after: (**A**) Removal of firing rate fluctuation from each neuron in SWS. (**B**) Estimating SWS cvPCs using up state periods only. (**C**) Subsampling the spikes from top half firing rate neurons to match the amount of spikes of bottom half. (**D**) Using solely putative pyramidal cells. [repeated measures ANOVA, multiple comparisons **p<0.05, **p<0.01, ***p<0.001]

**Fig. S 8.**
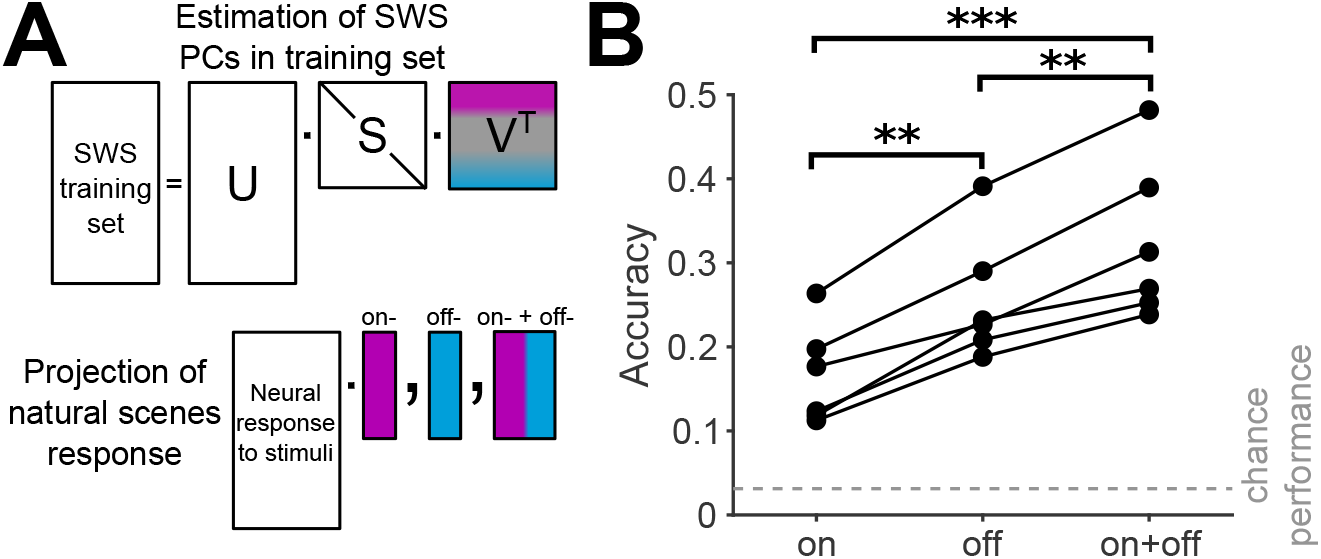
The on- and off-manifold subspaces encode complementary information about natural scene stimuli. (**A**) To test whether the on- and off-manifold subspaces encode redundant information, we tested the performance of a classifier in predicting stimulus identity from neural activity in the on-manifold subspace, the off-manifold subspace, or both. (**B**) Prediction performance is higher for both subspaces combined. [repeated measures ANOVA, multiple comparisons **p<0.01, ***p<0.001]

**Fig. S 9.**
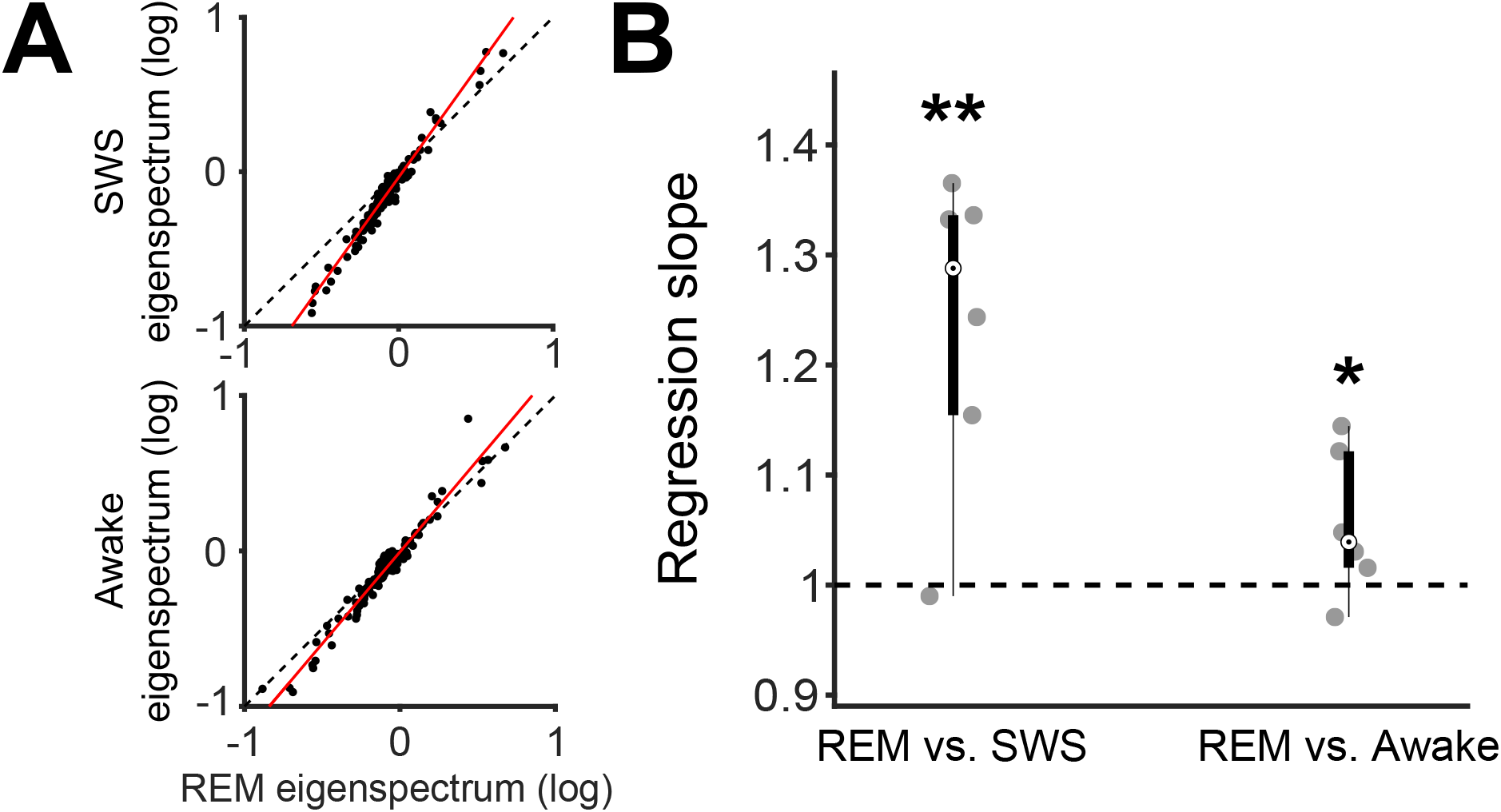
REM sleep is higher dimensional than SWS. (**A**) The eigenspectrum decays more slowly in REM, indicating higher-dimensional population activity than in SWS. (**B**) Summary of several recording sessions from V1 (SWS: two-tailed t-test, p = 0.0099, 1.24 ± 0.06, n = 6 recordings; Awake: p = 0.0955, 1.06 ± 0.03, n = 6 recordings; mean ± s.e.m.).

**Fig. S 10.**
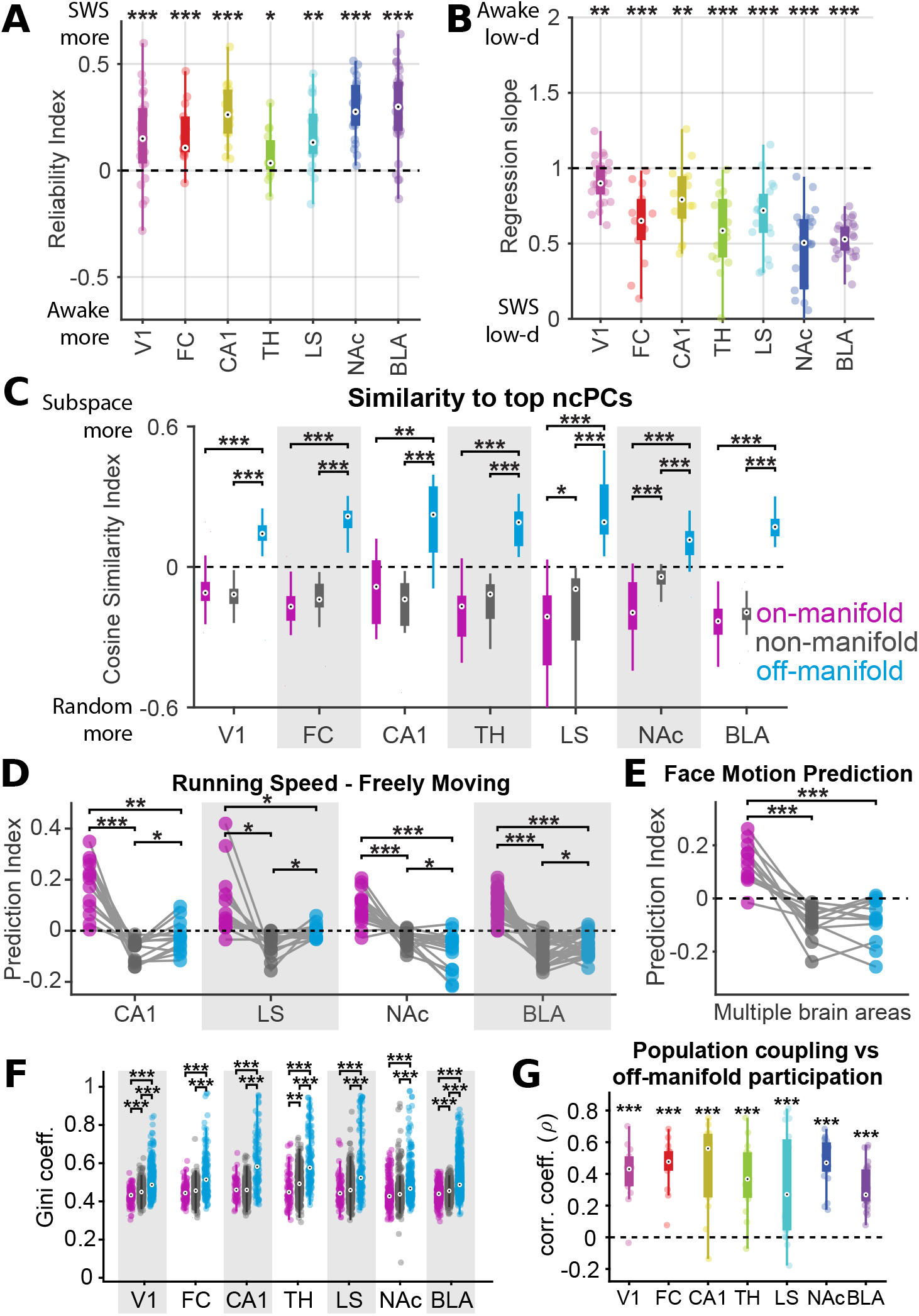
Main results found in V1 are replicated in multiple brain areas. (**A**) Population structure from SWS is more reliable than wakefulness for all brain areas tested (**B**) Eigenspectrum decay reveals that SWS is lower-dimensional than wakefulness in all brain areas tested. (**C**) Normalized contrastive PCA finds the dimensions maximally overrepresented in wakefulness; these dimensions are more aligned with the off-manifold subspace than the other subspaces. (**D**) During wakefulness, running speed encoding is on-manifold in CA1, LS, NAc and BLA datasets. (**E**) Neuropixels recordings from multiple brain areas have face motion encoding on-manifold but not non-/off-manifold. (**F**) For all datasets analyzed, off-manifold dimensions are sparser than on-/non-manifold dimensions. (**G**) Neurons participating in off-manifold coding are strongly coupled to the rest of the population. [*p<0.05,**p<0.01, ***p<0.001]

**Fig. S 11.**
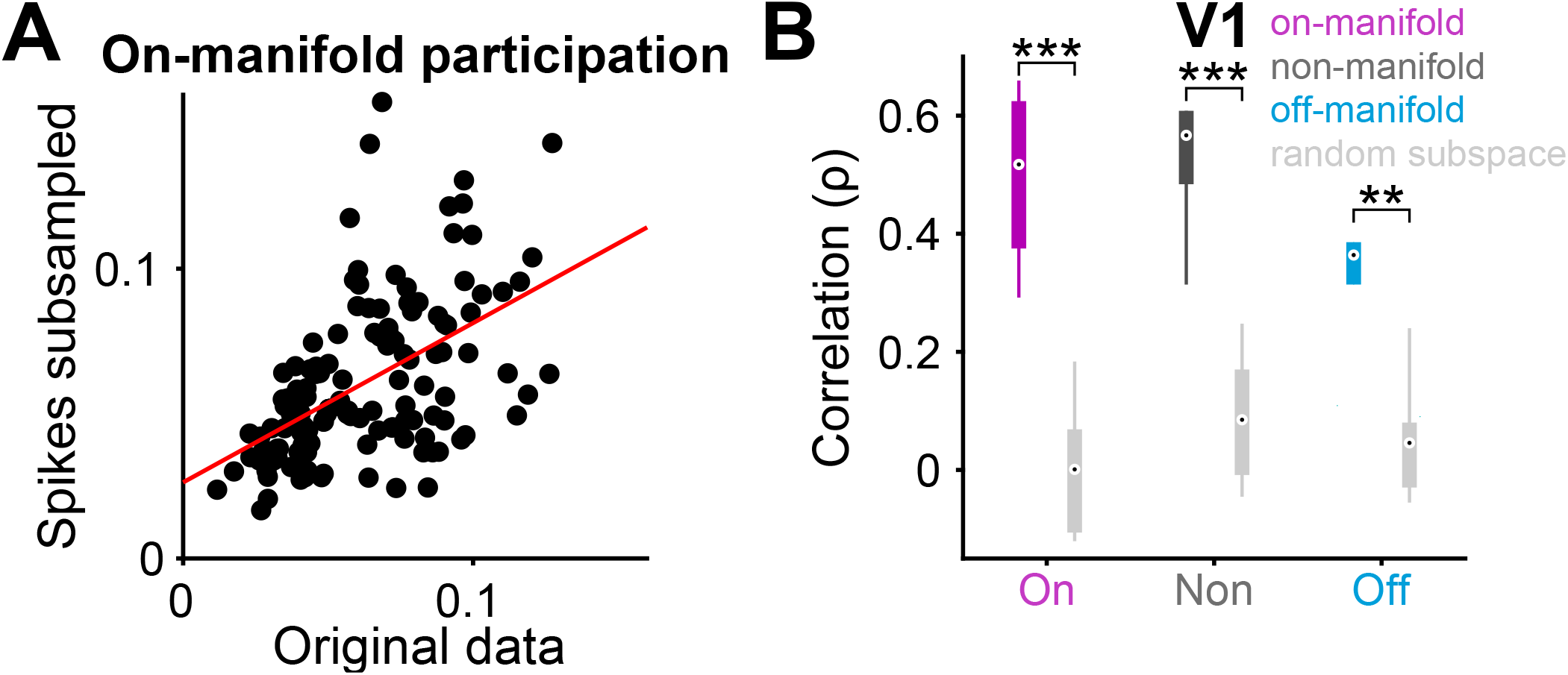
Subsampling spikes in high firing rate neurons does not drastically change subspace participation. (**A**) In an example session, the on-manifold participation of each neuron is similar before and after subsampling spike trains to control for firing rate. (**B**) Correlation coefficients for subspace participation in original and subsampled spike trains for all V1 recording sessions: on-manifold (magenta), non-manifold (dark gray), and off-manifold (cyan). Light gray boxplots are for randomly chosen vectors. Pearson correlation, on-manifold: r = 0.49 ± 0.04, non-manifold: r = 0.6 ± 0.06, off-manifold: r = 0.4 ± 0.08 [paired t-test, **p<0.01, ***p<0.001]

**Fig. S 12.**
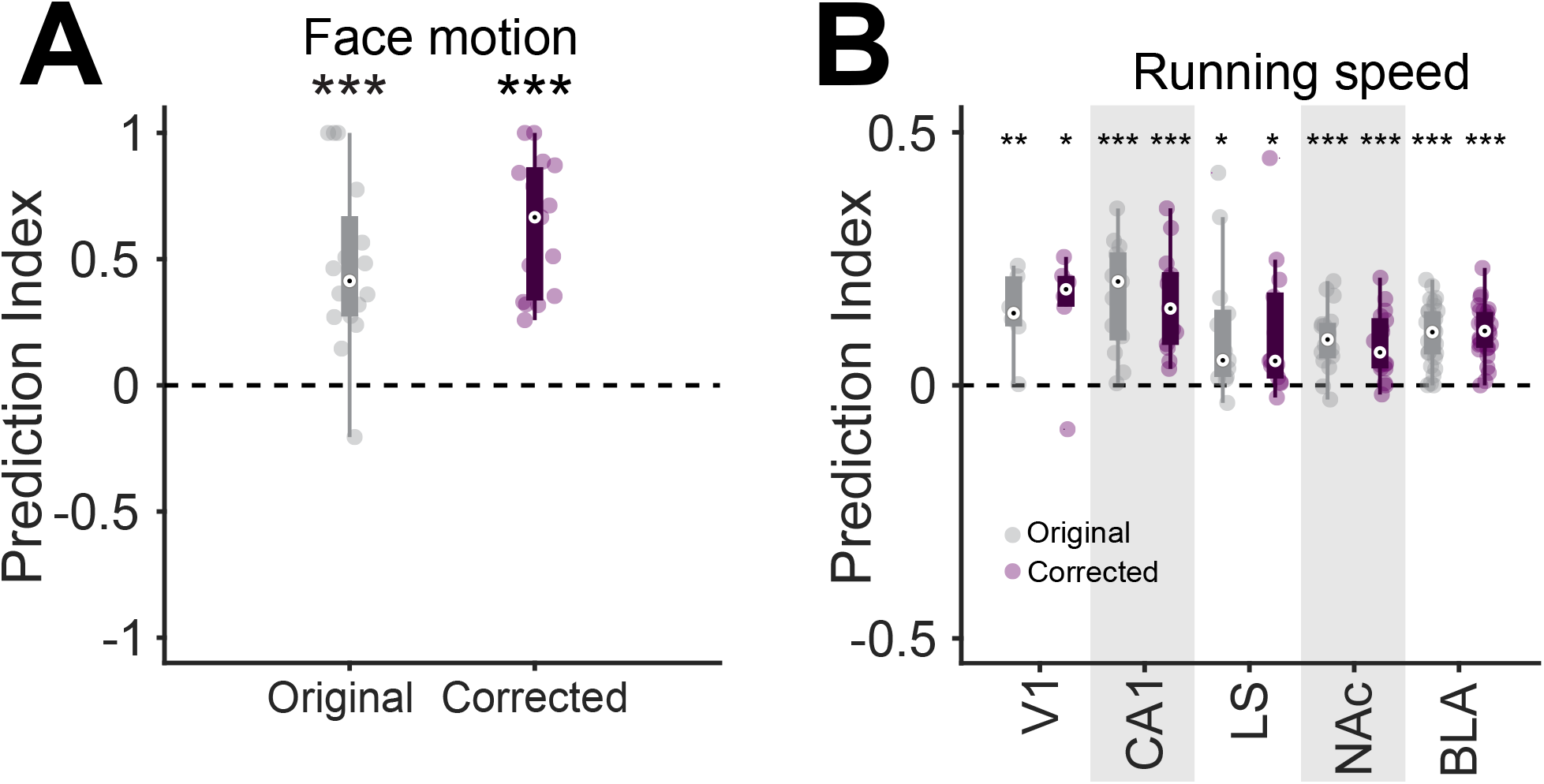
Controlling for firing rate fluctuations does not change on-manifold prediction of movements. (**A**) Removing overall firing rate fluctuation of each neuron does not change on-manifold predictability of face motion. Prediction index, original: 0.5 ± 0.09, corrected: 0.62 ± 0.07, mean ± s.e.m., n = 15 recordings of multiple brain areas. (**B**) Same as **A** but for running speed. Prediction index, original: 0.12 ± 0.01, corrected: 0.11 ± 0.01, mean ± s.e.m., n = 77 recordings. [two-tailed t-test, *p<0.05, **p<0.01, ***p<0.001]

**Fig. S 13.**
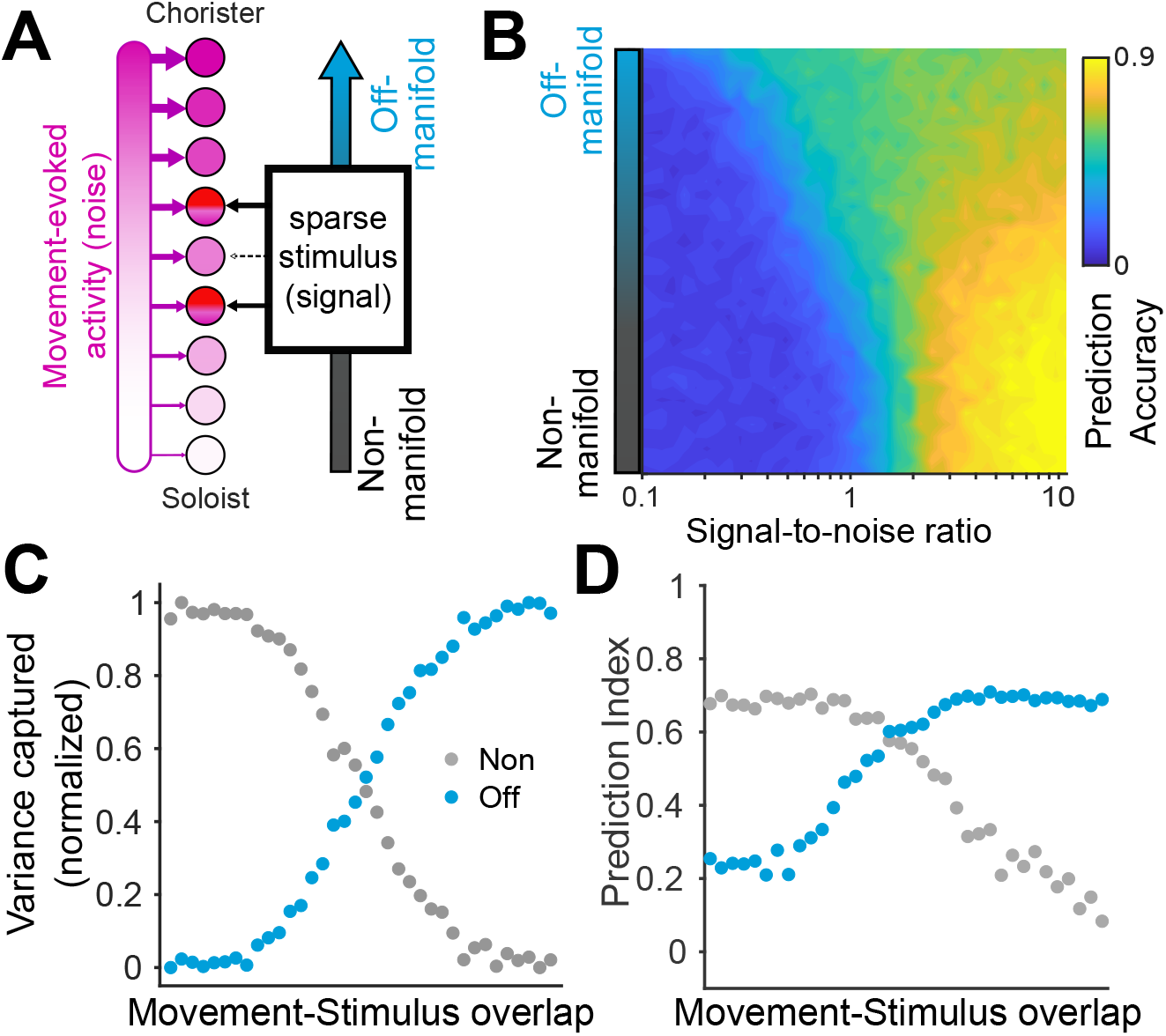
Stimulus-evoked activity in chorister neurons creates off-manifold encoding. (**A**) In our model, the stimulus provides inputs to a sparse subset of neurons that varies from low to high population coupling. (**B**) At low signal-to-noise ratios, off-manifold coding becomes advantageous. This is the same plot shown in Fig. 4F, but without normalization. (**C**) The proportion of stimulus-related variance captured by the non- and off-manifold subspaces (gray and cyan, respectively). As the sparse stimulus goes from overlapping with soloists to choristers, the coding of the stimulus switches from non-manifold to off-manifold. (**D**) The off-manifold prediction index increases as the sparse stimulus overlaps more with choristers than with soloists.

